# AXL-TBK1 driven nuclear AKT3 promotes metastasis

**DOI:** 10.1101/2022.01.17.476632

**Authors:** Emily N. Arner, Jill M. Westcott, Stefan Hinz, Crina Elena Tiron, Magnus Blø, Anja Mai, Reetta Virtakoivu, Natalie Phinney, Silje Nord, Kristina Y. Aguilera, Ali Rizvi, Jason E. Toombs, Tanner Reese, Vidal Fey, David Micklem, Gro Gausdal, Johanna Ivaska, James B. Lorens, Rolf A. Brekken

## Abstract

Epithelial-to-mesenchymal transition (EMT) contributes to tumor cell survival, immune evasion, migration, invasion, and therapy resistance. Across human cancer, tumors that are high grade, poorly differentiated, and have undergone EMT carry a worse prognosis with a higher likelihood of metastasis. AXL, a receptor tyrosine kinase, drives EMT and is implicated in tumor progression, metastasis, and therapy resistance in multiple cancer types including pancreatic cancer and breast cancer. TANK-binding kinase 1 (TBK1) is central to AXL-driven EMT yet, the mechanism of how TBK1 induces EMT remains unclear. Here, we report that AXL activation stimulates TBK1 binding and phosphorylation of AKT3. TBK1 activation of AKT3 drives binding and phosphorylation of slug/snail resulting in protection from proteasomal degradation and translocation of the complex into the nucleus. We show that nuclear translocation of AKT3 is required for AXL-driven EMT and metastasis. Congruently, nuclear AKT3 expression correlates with worse outcome in aggressive breast cancer. To advance AKT3 as a therapeutic target, an AKT3-isoform selective allosteric small molecule inhibitor, BGB214, was developed. BGB214 inhibits AKT3 nuclear translocation, EMT-TF stability, AKT3-mediated invasion of breast cancer cells and reduces tumor initiation in vivo. Our results suggest that AKT3 nuclear activity is an important feature of AXL-driven epithelial plasticity and that selective AKT3 inhibition represents a novel therapeutic avenue for treating aggressive cancer.

**Significance:** Nuclear AKT3 activity is an important feature of AXL-TBK1 driven EMT and metastasis, thus selective AKT3 targeting represents a novel approach to treat aggressive cancer.

## Introduction

Cancer metastasis, the leading cause of cancer mortality correlates with epithelial-to-mesenchymal transition (EMT) (1, 2). Metastasis of epithelial tumors, such as pancreatic cancer (PDA), requires cancer cells to escape epithelial nests, invade surrounding stroma, intravasate into blood or lymphatic vessels, survive circulation, and extravasate at a secondary site, where successful cells form micrometastases and eventually macrometastases (3). The escape of tumor cells from tumor cell nests encapsulated by a basement membrane can be facilitated by tumor cell epithelial plasticity, which results in epithelial tumor cells losing contact with the basement membrane and nearby cells while adopting mesenchymal-like features that enhance cell migration and invasion. While epithelial plasticity alters morphology and cell-cell contact it also enables tumor cell survival under stressful environmental conditions, such as chemotherapy and radiation (4–7).

In carcinomas, the manifestation of an EMT program is associated with tumor grade. High-grade cancer is characterized by a loss of normal tissue structure and architecture. High-grade tumors are often described as poorly differentiated, displaying tumor cells that have undergone full or partial EMT. In contrast, low-grade tumors that retain an epithelial phenotype are characterized as well-differentiated. Across human cancer, tumors that are high grade and poorly differentiated carry a worse prognosis with a high likelihood of metastasizing to distant organs (8). Understanding the molecular mechanisms underlying cellular plasticity and metastasis may reveal novel ways to target these programs for effective therapies.

Many signaling pathways can mediate tumor cell epithelial plasticity, including the receptor tyrosine kinase (RTK) AXL (9–11), elevated expression of which correlates with metastasis and resistance to therapy (9, 12). AXL is a member of the TAM (Tyro3, AXL, MerTK) family of RTKs (13) and is activated by its ligand, growth arrest-specific gene 6 (GAS6) to promote a variety of cellular processes, including epithelial plasticity, cell survival, proliferation and migration (9). We have previously shown that the serine threonine protein kinase TANK-binding kinase 1 (TBK1) promotes EMT downstream of AXL in PDA, providing insight into a novel function for TBK1 (14). While the precise mechanism of how TBK1 drives EMT has yet to be determined, previous work found that TBK1 can directly activate AKT (15).

AKT is a key regulator of many cellular phenotypes associated with cancer, including cell survival, proliferation, and metastasis (16). Activation of AKT can drive EMT via the induction of EMT transcription factors (EMT-TFs) including snail and slug, which transcriptionally repress E-cadherin and induce vimentin, twist1, MMP-2, and MMP-9 that promote tumor cell invasion (7, 17, 18). There are three mammalian AKT isoforms (AKT1, AKT2, and AKT3). While each isoform is encoded by distinct genes, there is ∼80% amino acid sequence identity and each isoform appears to be activated by similar mechanisms (19, 20). Although the function of AKT in general in cancer cell survival and growth has been well characterized, the contribution of different AKT isoforms has not been investigated as intensely and is often under appreciated. Based on a phosphoproteomics screen, AKT isoforms have specific expression patterns and serve different functions in cell signaling and cancer (21). Although it is the least studied isoform, AKT3 has been implicated in various aspects of EMT, including tumor progression, DNA damage repair response, and drug resistance (22–25).

Here we report a novel mechanism in which nuclear AKT3 is vital to AXL-TBK1 driven EMT by stabilizing the EMT transcription factors slug and snail. Additionally, we report the first AKT isoform specific small molecule inhibitor, BGB214, which is an AKT3-isoform selective allosteric small molecule inhibitor. BGB214 inhibits EMT-TF stability, AKT3-mediated invasion, and tumor initiation in vivo. Lastly, we show that AKT expression drives metastasis in vivo and nuclear AKT3 expression correlates with aggressive cancer. Our findings suggest that nuclear AKT3 activity is an important feature of AXL-driven epithelial plasticity and that selective AKT3 targeting represents a novel therapeutic avenue for treating aggressive cancer.

## Results

### AKT3 promotes EMT via TBK1

AXL activation promotes tumor cell migration and invasion (26). Consistent with this, *AXL* mRNA expression correlates with EMT and stem cell-related gene expression in breast cancer cell lines and patient breast carcinoma biopsies, but not normal breast tissue (Supplemental Figure 1A-C). Furthermore, IHC analysis of patient primary breast tumor biopsies revealed AXL protein expression correlates with expression of mesenchymal markers N-cadherin and twist2 (Supplemental Figure 1D). Interestingly, analysis of publicly available GEO RNA sequencing data of breast cancer cell lines showed that while *AXL* and *AKT3* correlate significantly, *AXL* and *AKT1* or *AKT2* do not (Figure 1A). Similar results were found by analyzing the correlation of AKT isoforms and AXL in human breast cancer using gene expression profiling interactive analysis (GEPIA) in invasive breast carcinoma (BRCA) from the TCGA database (Supplemental Figure 2A) (27). *AKT1* and *AKT2* showed no correlation with *AXL*, whereas *AKT3* correlated significantly with *AXL* expression in BRCA (*p*-value = 4.4×10^-110^, R = 0.61). In vitro, forced expression of slug in the epithelial breast line MCF10a (for cell line information see Supplemental Table 1) drives EMT and induces AXL and AKT3 expression, while AKT1 and AKT2 levels were not elevated (Supplemental Figure 2B). Additionally, when AXL was knocked down in these cells, AKT3 was no longer expressed (Supplemental Figure 2B), supporting the correlation between AXL and AKT3.

**Figure 1.**
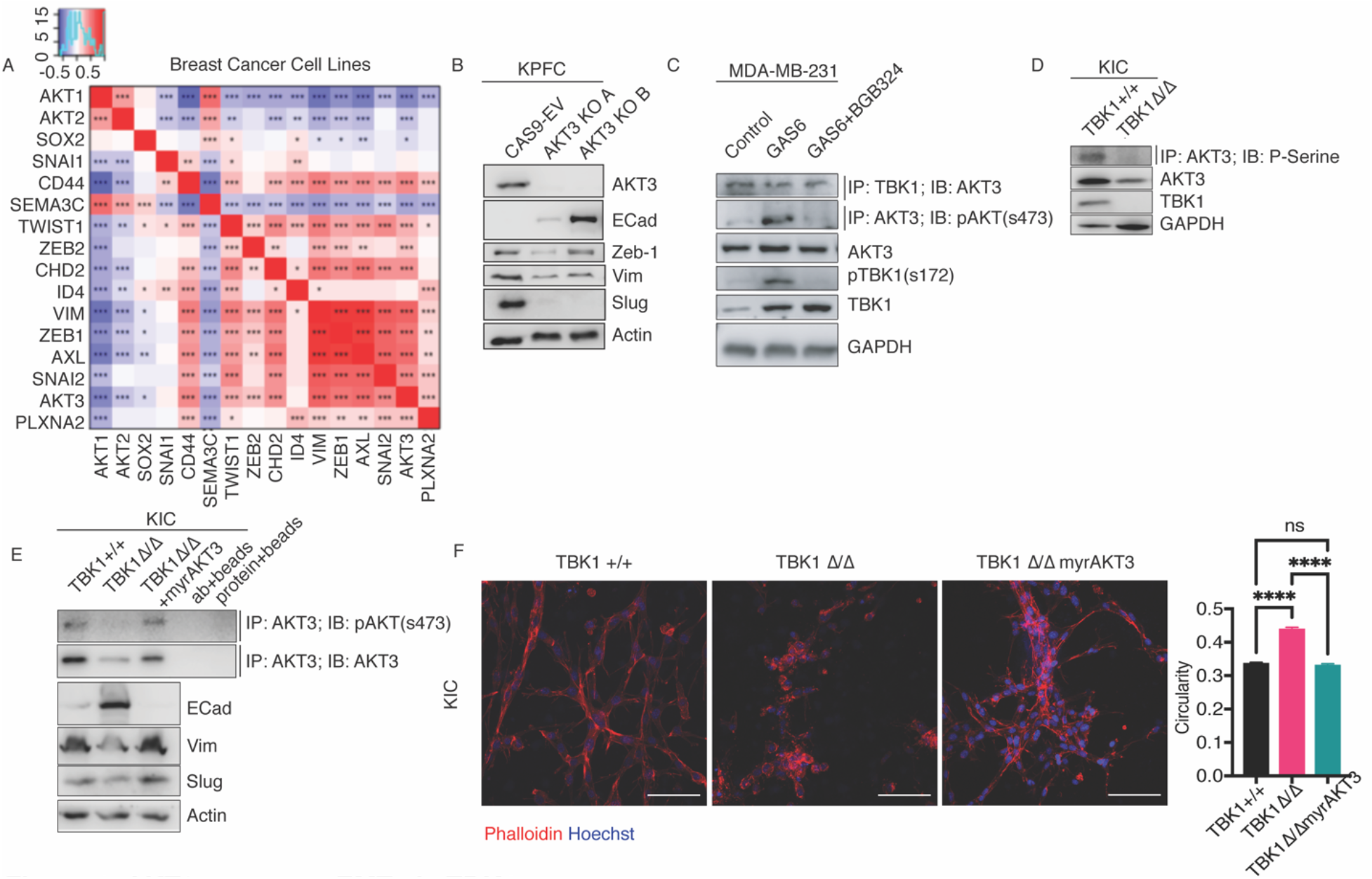
TBK1 activates AKT3 to promote EMT. **A)** Correlation of AKT1, AKT2, and AKT3 mRNA with EMT and stem cell associated genes in breast cancer cell lines. Positive correlation values are demarcated as red and negative correlation values are shown as blue (* p<0.045, ** p<0.009, *** p <2×10^-5^; Spearman’s correlation test). **B)** Western blot for the indicated target of *KPFC* PDA control Cas9-empty vector (CAS9-EV) or AKT3 CRISPR-mediated deletion (KO A, KO B). **C)** MDA-MB-231 cells were stimulated with DMSO, GAS6 (200 ng/ml) +/-2 μM BGB324. Immunoprecipitation of AKT3 was probed for pAKT(s473) and immunoprecipitation of TBK1 was probed for AKT3. Total lysates were probed for AKT3, pTBK1 (s172), TBK1 and GAPDH (loading control). **D)** Immunoprecipitation of AKT3 in *TBK1 WT* and *mutant* (*TBK1^Δ/Δ^*) *KIC* PDA cells probed for total phospho-serine. Total lysates were probed for AKT3, TBK1 and Actin (loading control). **E)** Immunoprecipitation of AKT3 in *TBK1^+/+^*, *TBK1^Δ/Δ^*, and *TBK1^Δ/Δ^* KIC PDA cells transduced with myrAKT3 (*TBK1^Δ/Δ^*-myrAKT3). AKT3 immunoprecipitation was probed for pAKT (s473) and total AKT3. Cell lysates were probed for E Cadherin, vimentin, slug and actin (loading control). Immunoprecipitation controls without protein or antibody are shown. **F)** *TBK1^+/+^*, *TBK1^Δ/Δ^*, and *TBK1^Δ/Δ^*-myrAKT3 cells were plated in collagen/matrigel and stained for Phalloidin (red) and Hoechst (blue). Z-stack (1 μm) images were taken by confocal microscopy at 20X magnification. Scale bar, 100 μm. Circularity of cells was calculated using ImageJ. n > 500 cells/condition. All statistics were done using one-way ANOVA; * p<0.05, ** p<0.01, *** p <0.001, **** p <0.0001. All representative results shown were reproduced in at least three independent experiments. representative results shown were reproduced in at least three independent experiments.

To investigate the function of AKT isoforms in EMT, MCF10a cells were treated for 4 days with TGFβ, a potent EMT inducer, after which each AKT isoform was immunoprecipitated and probed for phosphorylation (S473). We found that TGFβ-induced EMT results in phosphorylation of AKT3, but not AKT1 or AKT2, supporting that AKT3 is selectively associated with EMT (Supplemental Figure 2C). To further test the function of AKT3, CRISPR knockout of AKT3 was done in a primary pancreatic cancer cell line derived from *Kras^LSL−G12D/+^Trp53^fl/fl^Pdx1^Cre/+^* (*KPFC*), a genetically engineered mouse model (GEMM) of PDA. In the absence of AKT3, mesenchymal markers zeb-1, vimentin, and slug were reduced, while the epithelial marker E-Cadherin was increased in two different clones (AKT3 KO A and AKT3 KO B) compared to the Cas-9 empty vector (CAS9-EV) control, confirming the function of AKT3 in EMT (Figure 1B).

Given the correlation of AKT3 with AXL and EMT, we sought to determine if AKT3 contributes to AXL-mediated EMT. To mimic constitutively active AKT1 or 3, MCF10a cells were transduced with retroviral vectors expressing myristoylated AKT1 (myrAKT1) or myristoylated AKT3 (myrAKT3) and analyzed for changes associated with EMT (protein expression and morphology, Supplemental Figure 2D, E). Transduction of myrAKT1 did not alter cellular phenotype. However, myrAKT3 transduction resulted in robust changes in cell phenotype as well as EMT protein changes. Expression of AXL and mesenchymal markers vimentin and N-cadherin were elevated and the cells displayed a more invasive and mesenchymal-like morphology in 2D and 3D (embedded in matrigel), suggesting constitutively active AKT3 can drive EMT (Supplemental Figure 2D, E). To investigate if AKT3 is activated downstream of AXL, PANC1 cells were treated with DMSO, GAS6, or GAS6 and a neutralizing monoclonal anti-AXL antibody, tilvestamab. Probing for pAKT3 indicated that AKT3 can be activated in an AXL specific manner (Supplemental Figure 2F).

Our prior studies established that TBK1 promotes EMT downstream of AXL in PDA (14). Although the mechanism by which TBK1 drives EMT remains unclear, prior evidence shows that TBK1 can directly activate AKT (15, 16). Given our previous findings that AKT is activated downstream of AXL in a TBK1-dependent manner (14) we hypothesized that TBK1 binds to and activates AKT3 to drive EMT downstream of AXL. To test this, we treated MDA-MB-231 cells with DMSO, GAS6, or GAS6 plus BGB324 (R428; bemcentinib), a small molecule AXL kinase inhibitor (12, 28). Immunoprecipitation of TBK1 revealed that TBK1 binds to AKT3, and that AXL stimulation results in the phosphorylation of TBK1 and AKT3 (Figure 1C). Furthermore, BGB324 inhibited GAS6-induced activation of TBK1 and AKT3. To investigate TBK1-AKT3 interaction further, we used primary cell lines developed from GEMMs of pancreatic cancer, *TBK1^+/+^ KIC* (*Kras^LSL−G12D/+^ ; Cdkn2a^Lox/Lox^ ; Ptf1a^Cre/^*) or *TBK1-mutant* (*TBK1^Δ/Δ^*) *KIC* mice (14), which are deficient in TBK1 kinase activity. We found that AKT3 is phosphorylated in *TBK1^+/+^ KIC* cells but not in *TBK1^Δ/Δ^ KIC* cells (Figure 1D), supporting the hypothesis that TBK1 can activate AKT3. To investigate if TBK1 can directly bind to and activate AKT3 we performed an in vitro kinase activity assay with human recombinant TBK1 and AKT3 using cold ATP. Mass-spectrometry analysis confirmed that TBK1 directly phosphorylates AKT3 at serine 472 (Supplemental Figure 2G).

To investigate if AKT3 induces EMT downstream of TBK1, we transduced *KIC TBK1^Δ/Δ^* cells with myrAKT3 (*TBK1^Δ/Δ^*/myrAKT3) and found that myrAKT3 rescues expression of mesenchymal markers, vimentin and slug, and decreases the expression of E-cadherin (Figure 1E), demonstrating that myrAKT3 induces a mesenchymal-like phenotype in TBK1-mutant PDA cells. To evaluate if the protein changes seen in Figure 1E result in a phenotypic change, we cultured *TBK1^+/+^*, *TBK1^Δ/Δ^*, and *TBK1^Δ/Δ^*/myrAKT3 *KIC* cells in collagen/matrigel and found that *TBK1^+/+^* cells were invasive with elongated morphology while *TBK1^Δ/Δ^* cells were epithelial and less elongated (Figure 1F). Interestingly, *TBK1^Δ/Δ^*/myrAKT3 cells reverted to a mesenchymal-like morphology, similar to *TBK1^+/+^* cells, suggesting constitutively active AKT3 is sufficient to drive EMT, even in the absence of TBK1. These data support that AKT3 is downstream of TBK1 and is required for TBK1 driven EMT.

### AXL-TBK1 is required for AKT3 nuclear localization

It has been reported that while AKT1 and AKT2 are found in the cytoplasm and mitochondria, respectively, AKT3 is often found in the nucleus (29). We observed clear nuclear localization of AKT3 in MDA-MB-231 and MCF10A/slug cells (30) (Supplemental Figure 3A,B). Interestingly, AXL silencing in MCF10a/slug cells reduced AKT3 nuclear localization, suggesting that AXL mediates the nuclear localization of AKT3 (Supplemental Figure 3B). These data were validated further with cell fractionation experiments where AKT3 was primarily detected in nuclear fractions of MDA-MB-231 cells (Figure 2A) and HMLER cells transduced with myrAKT3 (Supplemental Figure 3C).

**Figure 2.**
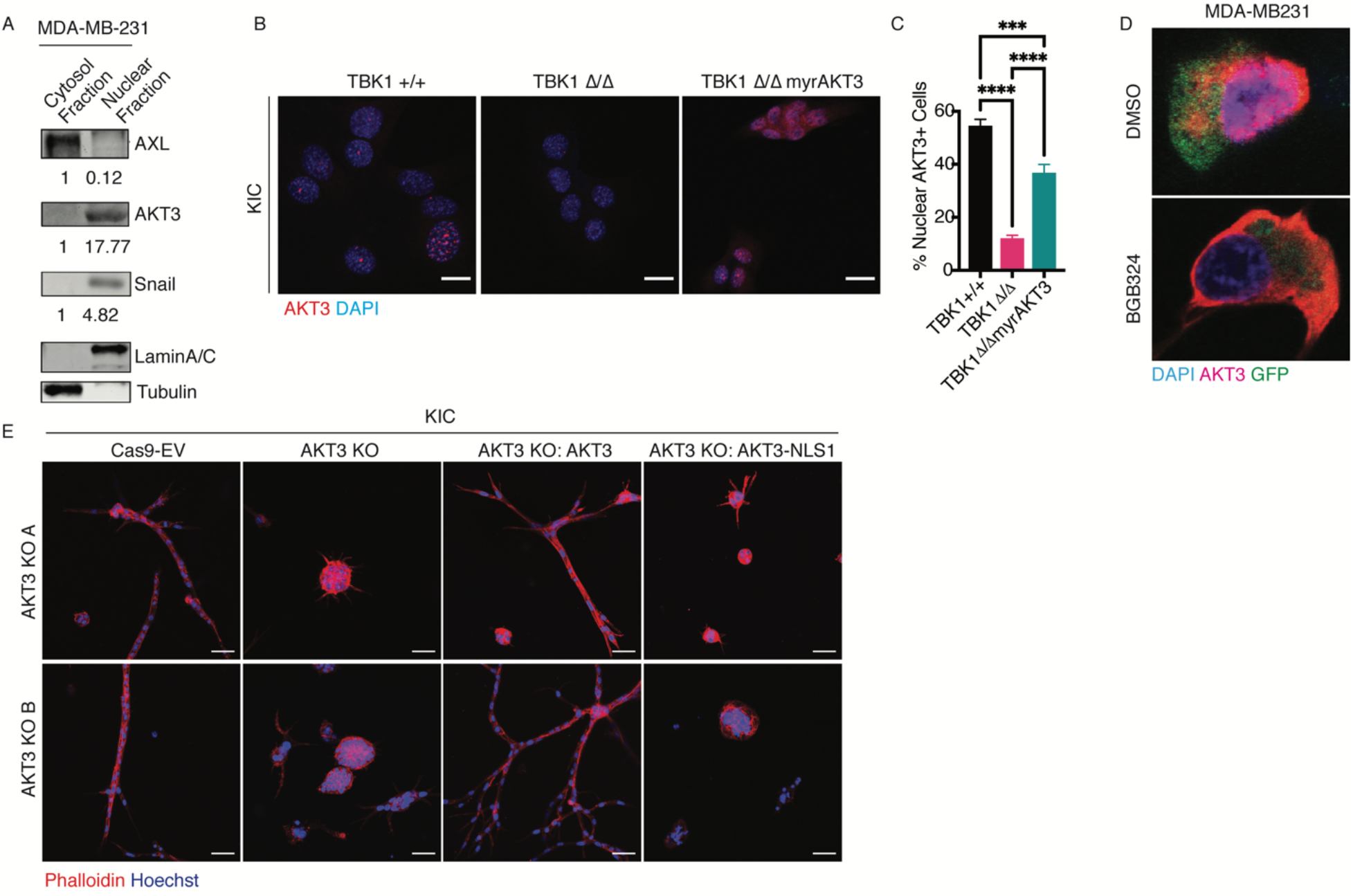
AXL-TBK1 is required for AKT3 nuclear localization. **A)** Western blot analysis of AXL, AKT3, snail, lamin A/C and tubulin in MDA-MB-231 nuclear and cytoplasmic fractions. Protein expression was quantified and normalized to protein in the cytosol. **B-C)** Immunofluorescence of AKT3 (red) and DAPI (blue) in *TBK1^+/+^*, *TBK1^Δ/Δ^*, and *TBK1^Δ/Δ^*-myrAKT3 *KIC* PDA cells. Percent of cells with nuclear AKT3 is graphed in (**C**). Scale bar, 20 μm. **D)** Immunofluorescence of MDA-MB-231/GFP cells treated with DMSO or BGB324. Cells are stained with AKT3 (red) and nuclei are stained with DAPI (blue). **E)** KPFC CAS9-EV, AKT3 KO, AKT3 KO cells transduced with AKT3, and AKT3 KO cells transduced with AKT3-NLS1 mutant plated in collagen/matrigel and stained with Phalloidin (red) and Hoechst (blue). Z-stack (1 μm) images were taken by confocal microscopy at 20X magnification. Cells were imaged at 20X using confocal microscopy. Scale bar, 50 μm. Two different AKT3 KO CRISPR clones are displayed. All representative results shown were reproduced in at least three independent experiments. All statistics were done use one-way ANOVA: ***, p <0.001; ****, p <0.0001.

To determine if TBK1 contributes to the nuclear localization of AKT3, immunofluorescence of AKT3 in *TBK1^+/+^*, *TBK1^Δ/Δ^*, and *TBK1^Δ/Δ^*/myrAKT3 *KIC* and MDA-MB-231 cells (Figure 2B, C) was performed. The percentage of cells with nuclear AKT3 was reduced ∼80% in the absence of functional TBK1 in *KIC* cells. This effect was partially rescued by myrAKT3, suggesting that AKT3 activation by TBK1 contributes to AKT3 nuclear localization. Furthermore, when MDA-MB-231/GFP cells were treated with BGB324 to inactivate AXL, thereby preventing TBK1 activation, AKT3 did not translocate to the nucleus (Figure 2D). To investigate how AXL affects nuclear localization of AKT3, MDA-MB-231 (Supplemental Figure 3D) and PANC1 (Supplemental Figure 3E) cells were treated with serum free media (SFM), GAS6, or GAS6 + BGB324 for 12 hrs. Immunocytochemistry for AKT3 in MDA-MB-231 cells demonstrated that AKT3 was nuclear localized in 15.9% of cells treated with serum free media (SFM) while GAS6 treatment resulted in 47.1% of cells showing nuclear AKT3 (Supplemental Figure 3D). In contrast, AXL inhibition with BGB324, decreased nuclear AKT3 to only 2.9% of cells, supporting that AXL stimulation induces the nuclear localization of AKT3. Similar effects were observed in PANC1 cells (Supplemental Figure 3E). To demonstrate that inhibition of AKT3 nuclear localization is not a general phenomenon associated with RTK inhibition, HMECs were treated with imatinib, an inhibitor of ABL/CKIT/PDGFR. BGB324 reduced nuclear AKT3 but the imatinib did not (Supplemental Figure 3F-G).

Proteins over 40 kDa must be actively transported through the nuclear membrane by importins, which recognize and bind nuclear location sequences (NLS) (31). We used a web-based NLS mapper (32) and identified a weak bipartite NLS in the AKT3 amino acid sequence (accession number: Q9Y243) located in a flexible linker region between the PH-domain and kinase domain (Supplemental Figure 4A). Based on these *in silico* findings we created two AKT3 mutant overexpression constructs: AKT3-NLS1 and AKT3-NLS2. AKT3-NLS1 carries two point-mutations (K141R and R142A) that alter the leucine rich NLS region to the sequence that resembles the linker area in AKT2 (Supplemental Figure 4A). For AKT3-NLS2, a 10 amino acid sequence flanking the NLS was replaced to mimic a longer part of the linker region as coded in AKT2. Wildtype AKT3 and the mutants were retrovirally delivered and expressed in HMLER cells (Supplemental Figure 4B). Immunocytochemical analyses showed clear subcellular localization differences between control, AKT3, AKT3-NLS1 and AKT3-NLS2 transfected cells (Supplemental Figure 4C). Wildtype AKT3 was predominantly (87%) nuclear localized; however, AKT3-NLS1 and AKT3-NLS2 mutants were largely restricted to the cytoplasm with 18% and 29% nuclear localization, respectively. Immunoprecipitation of AKT3 in HMLER lysates and probing with α-importin showed that the AKT3-NLS1 mutant had impaired interaction with α-importin (Supplemental Figure 4D).

To further investigate the contribution of nuclear AKT3 to EMT, *KPFC* AKT3 KO cells were transduced with wildtype AKT3 or AKT3-NLS1 and cultured in collagen/matrigel (Figure 2E). Evaluation of invasion revealed that while AKT3 KO resulted in reduced invasive phenotype compared to control, when wildtype AKT3 was rescued so was the invasive phenotype. However, when AKT3-NLS was transduced into the AKT3 KO cells, invasion was no longer rescued (Figure 2E), suggesting that nuclear AKT3 is necessary to drive EMT.

### Snail and slug are AXL-TBK1 dependent substrates of AKT3

EMT is orchestrated by a limited number of transcription factors, considered to be the ultimate inducers of EMT (EMT-TFs). These transcription factors include the zinc finger transcription repressors snail (*SNAI1*) and slug (*SNAI2*) (2, 33, 34). Previously, we found that the activation of TBK1 in AXL-driven metastasis drives the engagement of slug and snail (14). We used GEPIA analysis to evaluate the correlation between the mRNA expression levels of SNAI2 and the different *AKT* isoforms in BRCA (Supplemental Figure 5A). Indeed, while *AKT3* correlated with *SNAI2* expression, *AKT1* and *AKT2* did not.

Given the presence of AKT3 in the nucleus, we sought to determine if AKT3 interacts with EMT-TFs. Immunoprecipitation of AKT3 in *KIC* PDA cells revealed that AKT3 associated with snail (Figure 3A). Furthermore, this complex remained intact in *KIC* PDA lines only when TBK1 was functional (Figure 3A), suggesting TBK1 is required for the interaction between AKT3 and snail. When PANC1 cells were treated with SFM or GAS6 (PANC1 cells produce GAS6, therefore there is a baseline level of GAS6-AXL signaling in cells treated with SFM) for 12 hrs, snail was found to be in the nucleus and cytoplasm of the cells (Figure 3B). However, when AXL was inhibited with BGB324, snail translocation to the nucleus was significantly reduced, suggesting that the AXL-TBK1-AKT3 pathway is involved in snail/slug nuclear translocation. Similar results were found with slug in MDA-MB-231 cells (Supplemental Figure 5B). This phenomenon was confirmed using imaging flow cytometry (Amnis Imagestream®), which clearly showed an ∼80% reduction of cells that display nuclear slug after AXL inhibition (Figure 3C, D).

**Figure 3.**
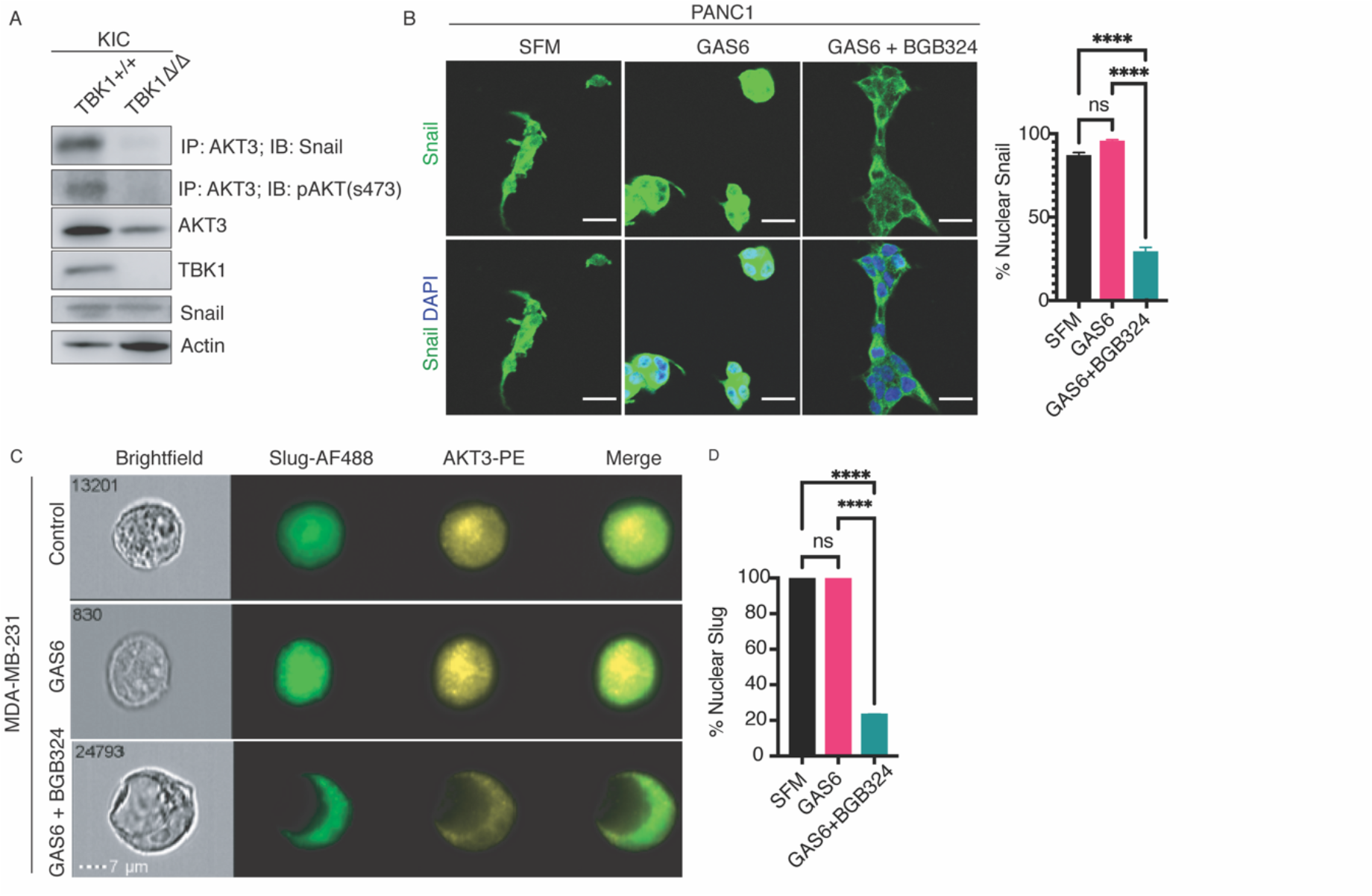
Snail/Slug is a TBK1-dependent substrate of AKT3. **A)** Immunoprecipitation of AKT3 in primary *TBK1 WT* and mutant (*TBK1^Δ/Δ^*) *KIC* cells were probed for snail and pAKT (s473). Lysates were probed for total snail, AKT3, TBK1, and Actin. **B)** Immunofluorescence of snail (green) and DAPI (blue) in PANC1 cells treated with serum free media (SFM), 200 ng/mL GAS6 +/-2 μM BGB324 for 12 hrs. Cells were imaged at 20X using confocal microscopy (scale bar, 20 μm) and nuclear snail was quantified, n>200 cells. **C**) Representative images from Imaging Flow cytometry (Amnis Imagestream®) of Slug-AF488 (green) and AKT3-PE (yellow) inMDA-MB-231 cells treated with 200 ng/mL GAS6 +/-2 μM BGB324 for 6 hrs. Scale bar, 7 μm. **D**) Nuclear localization of slug was quantified in each condition from (**C**); Control, n=549, GAS6, n=6781, GAS6 + BGB324, n=7730. Slug and AKT3 co-localization was quantified. All representative results shown were reproduced in at least three independent experiments. All statistics were done use one-way ANOVA: ****, p <0.0001.

### AXL activity stabilizes snail/slug via TBK1-AKT3

Given the interaction between AKT3 and snail (Figure 3A) and the strong effect of AKT3 expression on snail and slug expression (Figure 1B), we hypothesized that slug/snail activity is dependent on AKT3 providing a stabilizing effect on slug/snail protein. To test this hypothesis, MCF10a/slug cells were transfected with siAKT3. Even though slug was overexpressed to drive EMT in these cells (30), when AKT3 was not present vimentin and AXL expression were substantially reduced (Figure 4A), supporting the hypothesis that AKT3 is required for slug/snail EMT-inducing activity.

**Figure 4.**
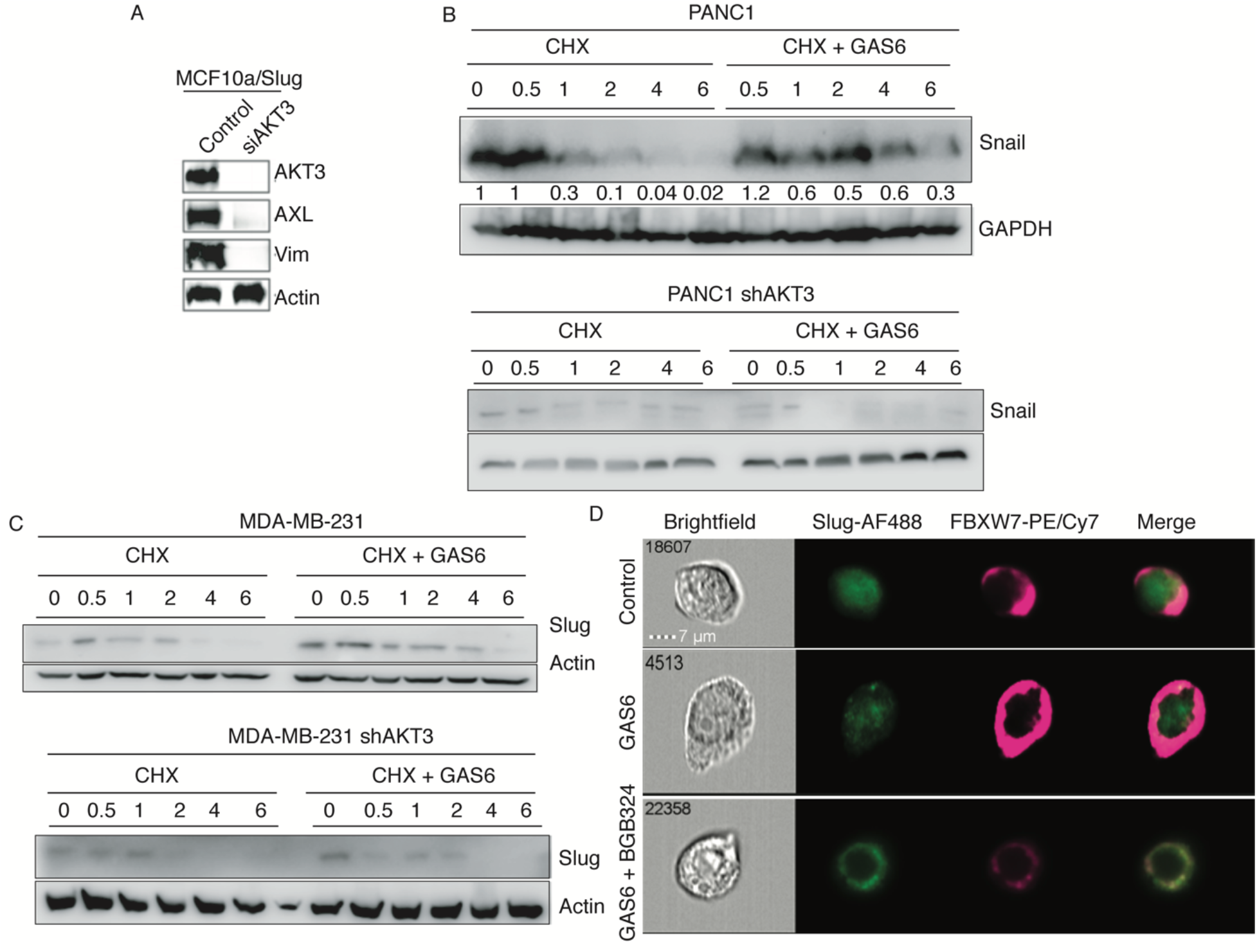
AXL activity stabilizes snail/slug via AKT3. **A**) MCF10a/Slug cells transfected with siAKT3 or control. Lysates were probed for AKT3, AXL, vimentin, and actin. **B,C**) PANC1 (**B**) and MDA-MB-231 (**C**) cells transduced with shAKT3 were treated with cycloheximide (CHX, 0.5 μg/mL) or CHX + GAS6 (200 ng/mL) and harvested at 30 min, 1, 2, 4, and 6 hrs of treatment. Lysates were probed for snail/slug and GAPDH/actin. **D**) Representative images from imaging flow cytometry (Amnis Imagestream®) of Slug-AF488 (green) and FBXW7-Pe/Cy7 (pink) in MDA-MB-231 cells untreated or treated with 200 ng/mL GAS6 +/-2 μM BGB324 for 6hrs. Scale bar, 7 μm. All representative results shown were reproduced in at least three independent experiments.

To determine if AXL-AKT3 activity influences the protein stability of snail/slug, we treated PANC1 (Figure 4B) and MDA-MB-231 (Figure 4C) cells with cycloheximide (CHX), a protein synthesis inhibitor, +/-GAS6 over a time course of 6 hrs. Consistent with previous findings (35) snail had a half-life of 1 hr when treated with CHX. Interestingly, when AXL was activated with GAS6, the half-life of snail was prolonged to 4 hrs, suggesting AXL activity stabilizes slug/snail protein. To determine AKT3 involvment, we repeated the experiment in cells transduced with shAKT3 and found that the addition of GAS6 no longer had a stabilizing effect on snail. AKT3 was similarly required for AXL-induced slug stabilty in MDA-MB-231 cells (Figure 4C). To determine whether snail protein is degraded by the proteasome or the lysosome, PANC1 cells were treated with a lysosome inhibitor (BafA1) or a proteasome inhibitor (MG-132) +/-GAS6 for 8 hrs (Supplemental Figure 5C). Although BafA1 had no effect on snail expression levels, when cells were treated with MG-132, there was a robust increase of snail protein, indicating snail is degraded via the proteasome.

The F-box E3 ubiquitin ligase FBWX7 has been implicated in the degradation of snail/slug in multiple cancers (36–38). Xiao and colleagues showed when FBXW7 was targeted with shRNA in two different lung cancer cell lines, the expression of snail increased markedly (36). This finding was recapitulated in ovarian cancer cells (37). To evaluate if AXL-AKT3 protects slug from FBXW7 and therefore degradation, we used Imaging flow cytometry (Amnis Imagestream®) of MDA-MB-231 cells and scored co-expression of FBXW7 and slug (Figure 4D). Interestingly, when MDA-MB-231 cells were treated with GAS6, FBXW7 and slug were rarely overlapping, but when AXL was inhibited using BGB324, overlap of the two proteins dramatically increased, suggesting that perhaps AXL-TBK1-AKT3 protects slug from FBXW7 mediated degradation.

### Selective targeting of AKT3 with a novel allosteric small molecule inhibitor inhibits metastasis

Several drugs targeting pan-AKT activity (e.g. GDC0068, AXD5363, MK-2206) are currently in various stages of clinical testing. However, many of these trials report toxicity such as hyperglycemia and hyperinsulinemia due to the essential functions of AKT1 and AKT2 in tissue homeostasis (39–42). An AKT3 selective inhibitor has the potential to overcome these issues. The similarity between AKT1, 2 and 3 in the kinase domain precludes selective kinase inhibition. However, an allosteric site located in a cleft between the PH domain and the kinase domain has been used to identify AKT1, AKT2 and AKT1/2-selective inhibitors (43–45). Sequence alignment around this allosteric site suggested that there are exploitable differences in this region (Figure 5A). A structural model produced by comparison of the crystal structures of AKT1 (45) and the AKT2 kinase domain suggested that a single amino acid deletion in AKT2 and AKT3 compared to AKT1 leads to a change in the path that the protein backbone follows, opening up a pocket at the front of the allosteric binding site (Figure 5B). This pocket is small in the case of AKT2 due to the protrusion of the large side chain of Arg269, but larger in AKT3 due to the presence of a glycine at this site. A series of novel allosteric small molecule inhibitors of AKT3 were developed (<u>WO/2016/102672</u>) with backbones that bind to the allosteric site via the right hand side of the molecule with the group on the left making a bend to access the additional space, causing the molecule to clash with AKT1 Lysine 268. One example of these is BGB214 (N-(5-(4-(1-aminocyclobutyl)phenyl)-4-phenylpyridin-2-yl)-2-((1r,4r)-4-(N-methylacetamido)cyclohexyl)acetamide), a potent and selective AKT3 inhibitor (Figure 5C). In biochemical assays using purified tag-free enzymes, BGB214 had an IC50 of 13 nM for AKT3 with approximately 1000-fold selectivity against AKT1 and >35-fold selectivity against AKT2 (Figure 5D).

**Figure 5.**
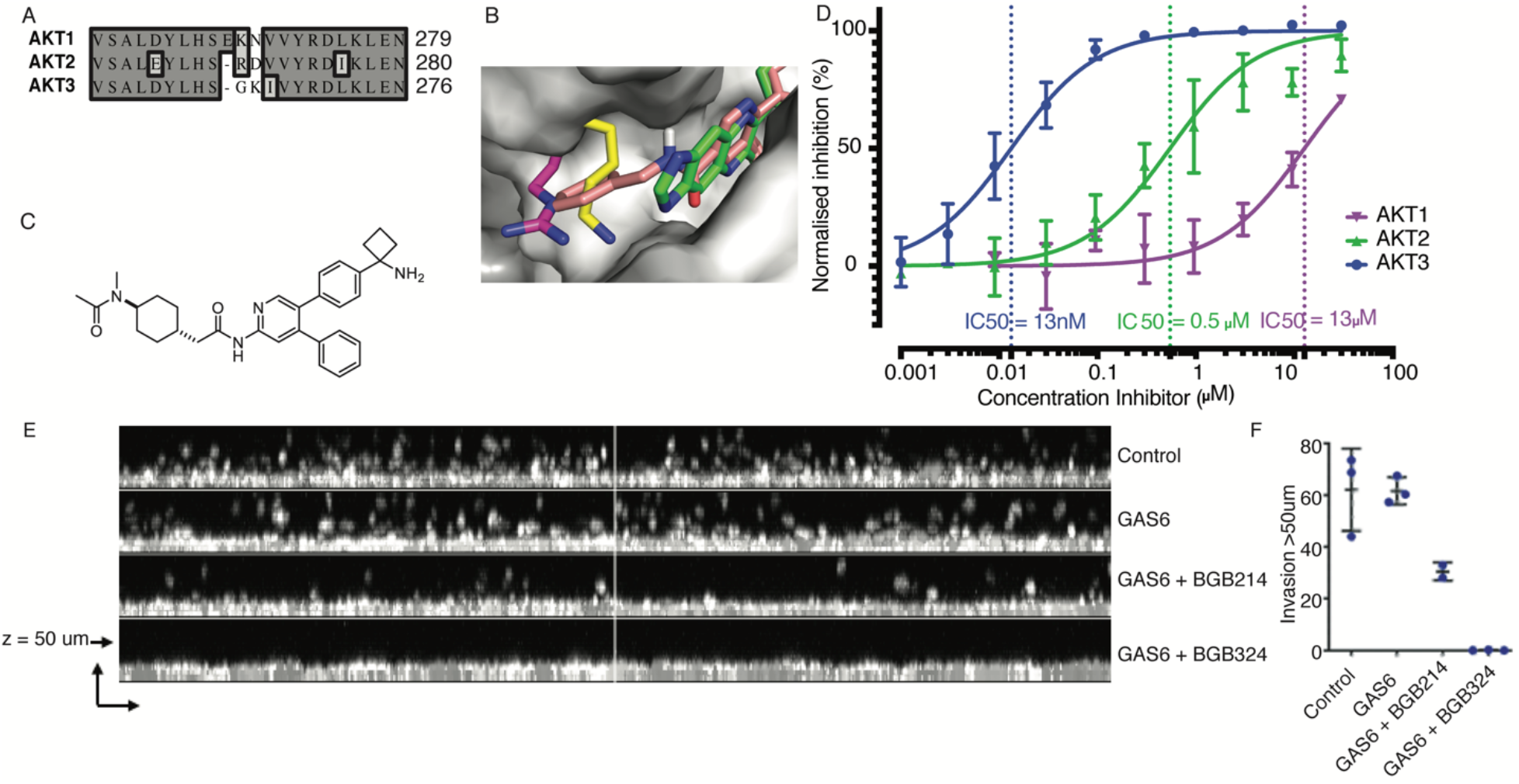
Efficacy of selective targeting of AKT3 with a novel allosteric small molecule inhibitor. **A**) Exploitable differences in sequence between AKT1, AKT2 and AKT3 around the allosteric site include a deletion in AKT2 and AKT3 compared to AKT1, implying that the backbone may follow a different path in these proteins. **B**) Surface view of the front of the allosteric binding site of AKT3, including bound allosteric inhibitor AKT VIII (green). Homology model of AKT3 based on crystal structures of AKT1 bound to AKT VIII (PDB 3o96) and AKT2 kinase domain (PDB1o6k). Side chains from the AKT1 crystal structure (Lys268, yellow) and the AKT2 crystal structure (Arg269, magenta) are superimposed, showing how they impinge on the space made available by the smaller glycine present at this location in AKT3 (Gly265). A molecule with similar structure to BGB214 (pink) docked at the allosteric site clashes with Lys268 of AKT1 (yellow). **C**) Structure of BGB214. **D**) Inhibition of AKT1, AKT2 and AKT3 enzymatic activity on GSK3α-derived *Ultra* U *light*TM-labelled crosstide substrate (n>3). **E**) MDA-MB-231 cells plated in collagen/matrigel and treated with GAS6 +/-3 μM BGB214 or 2 μM BGB324 for 48 hrs. Z-stack images were taken using confocal microscopy over 50 μm. **F**) Invasion greater than 50 μm was quantified.

To evaluate the efficacy of BGB214 to prevent aggressive cancer traits such as migration and 3D growth, MDA-MB-231 cells were plated in collagen/matrigel and treated with GAS6, GAS6 + BGB214, or GAS6 + BGB324 for 48 hrs (Figure 5E, F). Invasion over 50 μm was determined and quantified revealing that inhibition of AXL or AKT3 substantially reduced cell migration/invasion (Figure 5F). Similarly, in an organotypic 3D growth assay, BGB214 dose-dependently prevented MDA-MB-231 growth (Supplemental Figure 6A), but did not significantly affect cell growth in 2D proliferation assays (Supplemental Figure 6B, C). Interestingly, BGB214 inhibition in PANC1 cells resulted in decreased expression of total snail (Supplemental Figure 6D).

The specificity of BGB214 for pAKT3 was confirmed in a panel of cell lines in vitro and in vivo. HMLER-AKT3 or HMLER-GFP cells were treated with increasing concentrations of BGB214 (Supplemental Figure 6E). As HMLER-GFP cells have very low levels of AKT3 endogenously, pAKT levels were only reduced when AKT3 was overexpressed, indicating that BGB214 selectively inhibits AKT3. In addition, MCF10-DCIS subcutaneous tumors treated with 25 mg/kg BGB214 for 2-6 days specifically resulted in decreased pAKT3 with little effect on pAKT1 and pAKT2 (Supplemental Figure 6F).

To investigate the potential of BGB214 to prevent tumor initiation, HMLER cells transduced with control vector or AKT3 were pre-treated in vitro with BGB214 for 24 hours and then injected subcutaneously into NOD SCID mice at limiting dilutions (1×10^5^-1×10^6^ cells) and mice treated with BGB214 for 14 days (Supplemental Figure 6G). AKT3 inhibition by BGB214 significantly reduced the tumor initiation capacity of HMLER-AKT3 cells (Supplemental Figure 6G). The same reduction in tumor initiating capacity was observed following injection of HMLER-AKT3 cells without in vitro treatment with BGB214 preceding injection (Supplemental Figure 6H). We conclude that inhibition of AKT3 with the allosteric inhibitor BGB214 prevents AKT3 mediated tumorigenic features such as invasion, 3D growth, EMT transcription factor stability and tumor initiation.

### AKT3 expression is associated with poorly differentiated tumors and increased metastasis

To assess the biologic consequence of AKT3, control *KPFC* cells (CAS9-EV), AKT3 KO KPFC cells (AKT3 KO), or AKT3 KO KPFC cells transduced with AKT3 (Rescue) were injected orthotopically into the pancreas of C57BL/6J mice (Figure 6). Primary tumor and metastastic burden was evaluated 19 days post injection. Although tumor weight did not differ significantly between the three groups (Figure 6B), gross metastatic burden was reduced in tumors lacking AKT3 (Figure 6A). H&E analysis as well as CK19 (a PDA tumor cell marker, (46) IHC confirmed significantly reduced metastasis to livers of AKT3 KO tumor-bearing mice (Figure 6C-D). Consistent with this observation, tumors lacking AKT3 were more well-differentiated and expressed higher levels of E-Cadherin and lower levels of vimentin (Figure 6E). The expression of E-Cadherin and vimentin in vivo was consistent with the expression of these proteins in vitro (Figure 6F). Importantly, we observed that the expression level of AKT3 also correlated with the number of gross metastases (Figure 6A, F).

**Figure 6.**
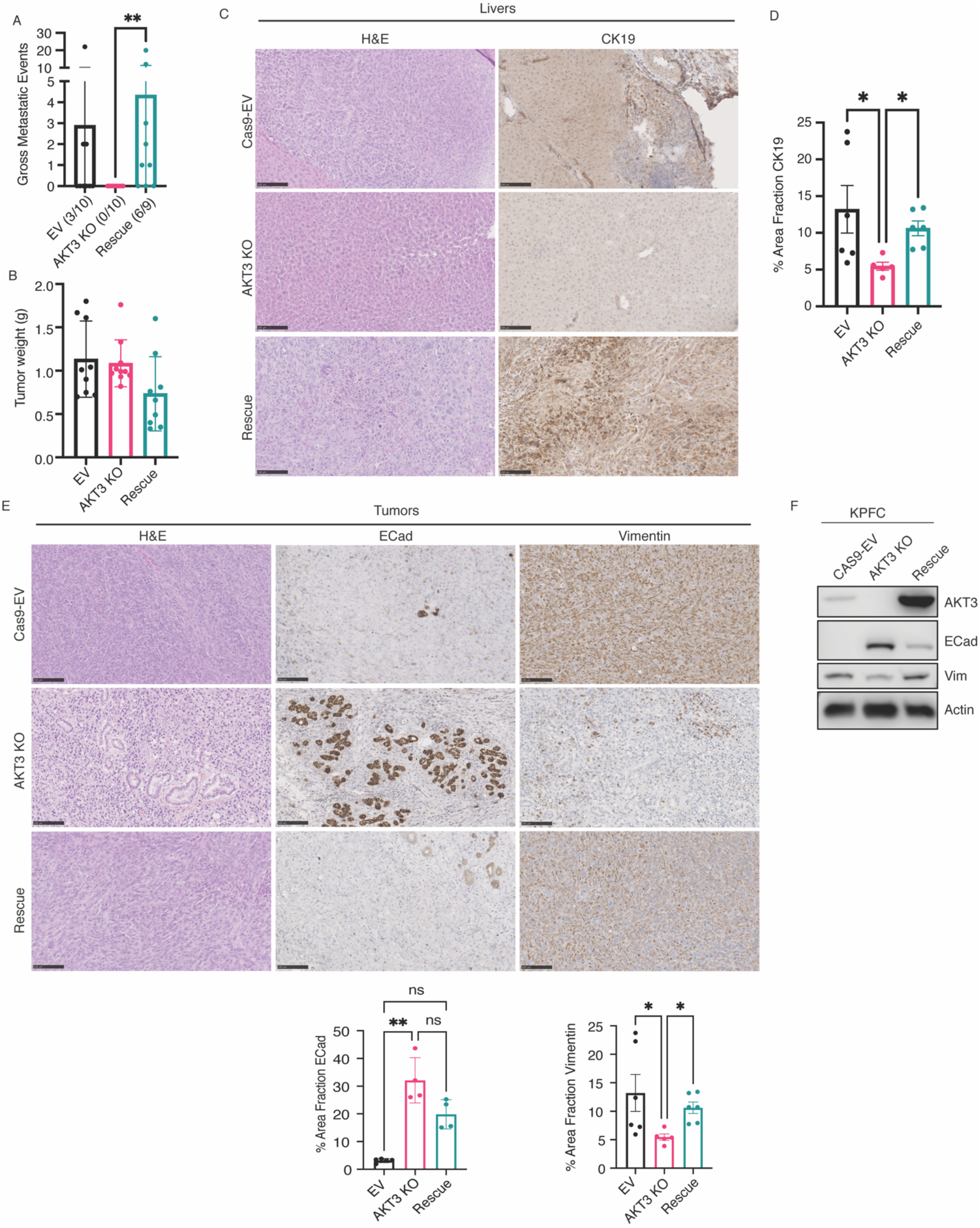
AKT3 expression is associated with poorly differentiated tumors and increased metastasis. C57BL/6J mice were injected orthotopically with 250,000 *KPFC* PDA cells (CAS9-EV, AKT3 KO, or Rescue). **A)** Gross metastases (CAS9-EV: 3/9; AKT3 KO: 0/10; Rescue: 6/9) and **B)** primary tumor weight was evaluated 19 days after tumor cell injection. **C-D)** Representative images of H&Es and CK19 IHC on livers. CK19 reactivity was quantified as percent of total liver area. **E)** Representative images of tumors stained using H&E and IHC for E-Cadherin and vimentin. Percent area of Ecad and vimentin was quantified. **F)** Western blot of KPFC CAS9-EV, AKT3 KO, and Rescue cells. Cells were lysed and probed for AKT3, ECad, vimentin, and actin (loading control). All statistics were done using one-way ANOVA: *, p<0.05; **, p<0.01.

### Nuclear AKT3 is associated with aggressive cancer and worse survival in patients

We next sought to evaluate the importance of nuclear AKT3 in cancer patients. To assess the location of AKT3 in pancreatic tumors from patients, IHC for AKT3 and AXL (Figure 7A) demonstrated that AXL^+^ tumors displayed single cells outside epithelial ducts that expressed nuclear AKT3. However, in AXL^-^ tumors, AKT3 was cytoplasmic, supporting our findings that AXL is associated with nuclear localization of AKT3 and this localization results in a less differentiated (more mesenchymal-like) tumor cell phenotype (Figure 7A).

**Figure 7.**
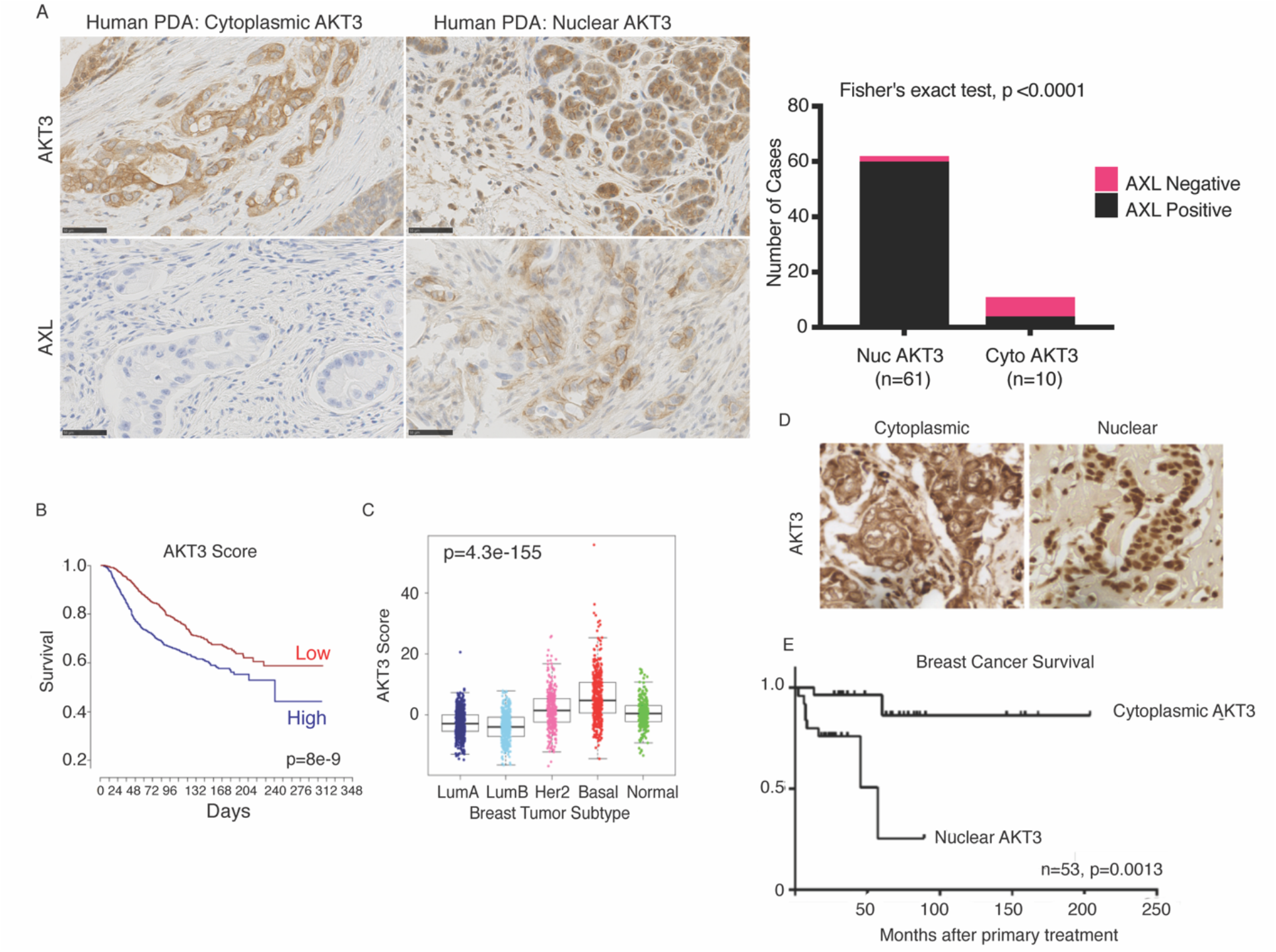
Nuclear AKT3 is associated with aggressive cancer and worse survival in patients. Representative images of IHC staining for AKT3 and AXL in human PDA (n=71). Fisher’s exact contingency test was used to calculate if there was a correlation between nuclear AKT3 and AXL positive expression within the tumor. **B**) Kaplan Meier plot indicating that high AKT3 induced expression (AKT3 score) correlated with worse outcome (p=8e-9) based on the METABRIC database. **C**) Significant different distribution of the AKT3 score between the PAM50 subtypes (p=4.3e-115, Kruskal–Wallis test); ER negative tumors were in general enriched for the AKT3 score, particularly Basal-like tumors. **D**) Representative images of IHC staining for AKT3 in human breast cancer samples in (**E**). AKT3 localization predominantly cytoplasmic (left) or nuclear (right). **E**) Survival analyses of 53 breast cancer patients based on nuclear or cytoplasmic AKT3 localization (p=0.0013 Log rank (Mantel-Cox) test).

To assess the effect of AKT3 expression in breast epithelial cells, we retrovirally overexpressed AKT3 in MCF10A cells and compared the mRNA expression pattern via RNA sequencing with MCF10A cells transduced with GFP control vector. We found 46 differentially expressed (DE) genes (FC≥2, FDR<0.05) (Supplemental Table 2). The DE genes and their directionality were used to calculate an “AKT3 score” which was then mapped against probes in the Metabric database, which is composed of gene expression patterns from 1980 breast cancer patients. The patients were divided into two groups depending upon if the AKT3 score was above or below the mean. Plotting the AKT3 score against patient survival indicates that a high AKT3 score correlates with a significantly worse overall outcome (KM, p=8e-9) (Figure 7B). Further, we found a significantly different distribution between breast cancer subtypes (based on PAM50 intrinsic subtypes) and AKT3 scores (p=4.3×10^-155^, Kruskal–Wallis test) (Figure 7C). ER negative tumors are in general enriched for the AKT3 score, particularly basal-like tumors (Figure 7C). That high levels of AKT3 associated gene expression correlates with more aggressive forms of breast cancer, worse overall outcome, and a higher hazard ratio is consistent with previous reports of AKT3 high copy number alterations in TNBC patients (47, 48) and reports that AKT3 expression is associated with higher grade breast cancer tumors (49). To validate our previous findings, we sought to determine if nuclear AKT3 was associated with worse overall survival. To evaluate this, we performed IHC for AKT3 in clinical breast cancer samples (Figure 7D, E). Grouping patients based on AKT3 subcellular localization revealed that nuclear AKT3 predicted a worse overall outcome (n=53 patients, p=0.0013 Log-rank test) in this cohort of patients. Together, these results suggest that nuclear AKT3 may be a therapeutic target that avoids toxicity associated with pan-AKT inhibition and a biomarker for worse overall survival and aggressive cancers.

## Discussion

We report that AXL activation by its ligand GAS6 leads to the stimulation of TBK1 and subsequent selective activation of the AKT isoform, AKT3. Activation of AKT3 drives the binding of AKT3 to its substrate slug/snail, and translocation into the nucleus. The binding of AKT3 to slug/snail also protects the EMT-TFs from proteasomal degradation, potentially by preventing ubiquitinylation by FBWX7 (Figure 8). These results highlight the function of AKT3 in EMT and its potential value as a therapeutic target, inhibition of which could enhance sensitivity to standard therapy.

**Figure 8.**
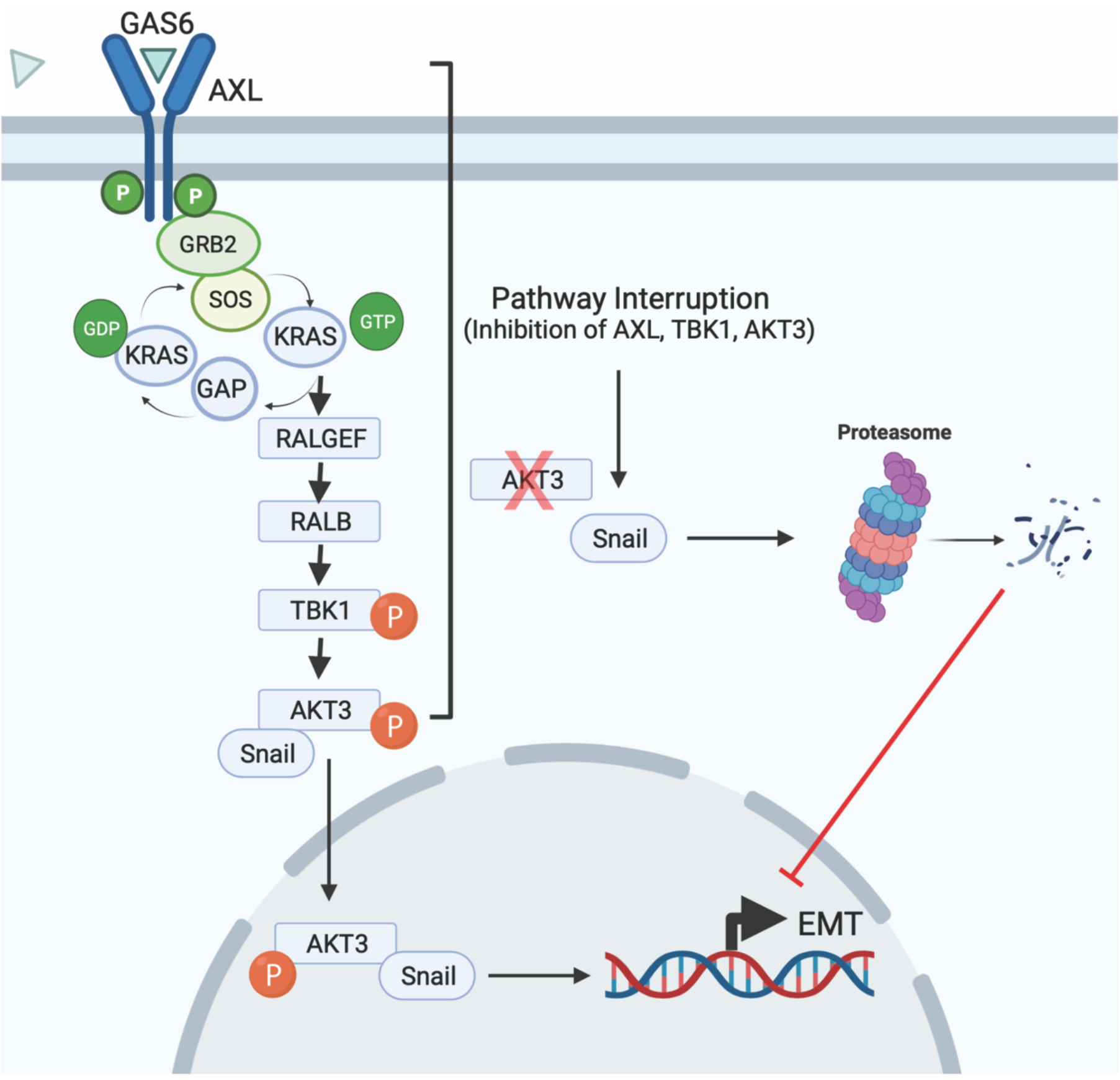
AXL-TBK1 driven nuclear AKT3 stabilizes slug and snail to promote EMT. AXL activation by its ligand GAS6 leads to the stimulation of TBK1 and subsequent activation of AKT3. Activation of AKT3 drives the binding of AKT3 to slug/snail, where they are translocated into the nucleus. The binding of AKT3 to slug/snail protects the EMT-TFs from proteasomal degradation. When this pathway is interrupted and AKT3 is not activated, AKT3 can no longer bind to slug/snail thus leading to proteasomal degradation of the EMT-TFs and a decrease in EMT.

AKT activation is linked to fundamental signaling pathways underlying cancer development and progression. Many investigations have focused on the function of AKT1, AKT2, or panAKT, but have largely ignored AKT3. This may be because AKT3 is the least expressed of the three isoforms (50) and prior results on the function of AKT3 in tumorigenesis are inconsistent (22, 51–55). Regardless, a few reports have suggested that AKT3 contributes to cancer progression, including breast cancer (56–59). We predict that the inconsistency of AKT3 studies may be due to differential genetic contexts of the studies as well as to the fact that AKT3 can be expressed as two alternatively spliced variants, one which lacks S472 (60). In the study by Suyama et al., overexpression of the AKT3 variant lacking S472 was associated with improved overall survival and reduced lung metastasis in preclinical models of breast cancer, whereas when AKT3 had the S472 phosphorylation site they saw increased tumorigenesis (60). This is consistent with our findings that phosphorylation on serine 472 via TBK1 is needed for AKT3 nuclear localization to promote EMT and metastasis.

Our previous studies have shown an increase in other EMT-TFs, such as ZEB1, downstream of AXL-TBK1 (14). Further studies are needed to evaluate if the AXL-TBK1-AKT3 signaling cascade only influences protein expression of snail/slug or multiple EMT-TFs. Additionally, in our study, we only interrogated this pathway in the context of AXL and AXL stimulation. It is possible that this mechanism may only be relevant in cell lines that contain high levels of AXL, which is supported by our finding that AXL is expressed in human PDAC tumors that display the nuclear localization of AKT3. Further studies are needed to understand if other RTKs can activate TBK1-AKT3 to stabilize snail/slug.

In our study we find that the binding of AKT3 to slug/snail protects the EMT-TFs from proteasomal degradation, although more studies are needed to determine if FBXW7 is required for the degradation of slug/snail. It is possible that AKT3 does not directly stabilize EMT-TFs, but perhaps other proteins such as deubiquitinating enzymes (DUBs) promote the stability of these EMT-TFs in an AXL-TBK1 dependent manner. For example, the DUB USP10 has been shown to promote the stability of slug and snail in breast, ovarian, and lung cancer cell lines (61). Other potential candidate proteins that might be involved in the degradation of snail are the F-box ligases, of which FBX15 and FBXO11 have been shown to ubiquitinate and support the degradation of Snail (62). Another protein that has been implicated in regulating the expression of slug and AXL is the transcription factor ΔNp63a, which has been shown to drive the migration of basal breast cancer cells in part through elevation of the expression of AXL and slug, as well as miR-205 to silence ZEB1/2 (63). ΔNp63a drives breast cancer invasion by selectively engaging certain proponents of the EMT program while still promoting the retention of epithelial characteristics to drive collective migration. Further studies are needed to evaluate the exact pathway by which slug/snail is degraded in an AXL-TBK1-AKT3 dependent manner.

These studies do not rule out that AKT3 affects other cell types such as macrophages; this is relevant given the function of TBK1 in innate immune signaling and the STING pathway. In fact, it has been reported that 7-DHC, a cholesterol precursor, regulates type I interferon production via AKT3 activation, where AKT3 directly binds and phosphorylates IRF3 on S385 (64). Additionally, AKT3 (pS473) in macrophages has been shown to promote migration, proliferation, wound healing, and collagen organization (65). Interestingly, a recent study showed that AKT3 phosphorylated RNA processing proteins that regulate the alternative splicing of fibroblast growth factor receptors (FGFR), consistent with an importance of nuclear targeting of AKT3 in EMT maintenance (21).

In summary, our data support that nuclear AKT3 has utility as a potential biomarker for aggressive cancers that express AXL, and that AKT3 is a specific mediator of EMT signaling downstream of AXL. Additionally, as there are ongoing clinical trials targeting AXL in multiple cancer types, analyses of these tumors for AXL expression and AKT3 localization after treatment may provide clinicians with a much-needed read-out for treatment efficacy. Lastly, we propose that selective inhibition of AKT3 may represent a novel therapeutic avenue for treating aggressive and recurrent cancer that avoids toxicity associated with pan-AKT inhibition.

## Materials and Methods

### Reagents

The following antibodies were used for immunoblotting (IB) at 1:1,000 unless otherwise stated: Anti-AXL (8661S, Cell Signaling, IHC 1:500), anti-phospho AXL y702 (5724, Cell Signaling), anti-phosphoserine (AB1603, Millipore), anti-TBK1 (3013S, Cell signaling), anti-TBK1 (ab40676, Abcam, IF 1:250), anti-TBK1 (NB100-56705AF647, Novus, FC 1:100), anti-pTBK1 s172 (5483S, Cell Signaling), anti-SNAIL (3879, Cell Signaling), anti-SLUG (9585, Cell Signaling), anti-SLUG-AF488 (NBP2-74235AF488, Novus, FC 1:100), anti-E-cadherin (clone 24E10, 3195S, Cell Signaling; IB 1:1000; IHC 1:400), anti-N-cadherin (14215S, Cell Signaling), anti-B-Catenin (8480, Cell Signaling), mouse anti-human Twist (Twist2C1a, Abcam, IHC), α-actin (A2066, Sigma, 1:2000), anti-α-tubulin (T6199, Sigma), anti-Vimentin (5741, Cell Signaling; IB 1:1000; IHC 1:400), anti-CK19 (ab52625, abcam; IHC 1:1000), anti-AKT1 (2967, Cell Signaling), anti-AKT2 (3063, Cell Signaling), anti-AKT3 (1586912, Millipore, IHC), anti-AKT3 (14982, Cell Signaling, IP, IB, and IF 1:250), anti-AKT3 (HPA026441, Sigma, IHC 1:200), anti-AKT3-PE (NBP2-71528PE, Novus, IF 1:250, FC 1:100), anti-pAKT (Ser473) (2971, Cell Signaling), anti-Lamin A/C IgG2b (Santa Cruz, sc-7292), anti-AKT (Cell signaling Technology, 9272), anti-Importin α (I1784, Sigma), anti-GAPDH (2118, Cell Signaling), anti-Phalloidin-AF546 (A22283, Invitrogen, IF 1:500), Hoechst 33342 (IF, 1:2000) and anti-FBXW7-PECY7 (NBP2-50403PECY7, Novus, FC 1:100). The following reagents were purchased from Sigma: Cycloheximide (01810-1g), BafA1 (B1793-2UG), MG-132 (474787-10MG). The CRU5-IRES-GFP retroviral vectors for expression of hSNAIL, hSLUG, myrAKT1, myrAKT3, AKT3, shLuc, and shAXL (RFP) were prepared as described (11). CRU5-IRES-GFP retroviral vectors for expression of AKT3-NLS and AKT3-NLS were generated by site-directed mutagenesis (Quik change #2200519). CRU5-IRES-GFP Luciferase AKT3-Luciferase and AKT3-NLS1-Luciferase were generated by cloning. All vectors were confirmed by DNA sequencing. Lentiviral shRNA constructs against human AKT1, AKT2, AKT3, and TBK1 were purchased from Dharmacon (TBK1, RHS3979-201735457, clone ID: TRCN0000003184)(AKT1, RHS3979-201768650, clone ID: TRCN0000039797)(AKT2, RHS3979-201732837, TRC00000005630)(AKT3, RHS3979-201733886, TRCN0000001612). Retroviral production and infections were conducted using Phoenix A retroviral packaging cells as described (66). Lentiviral production and infection were conducing using HEK293T cells as previously described (14). Human Gas6 from conditioned media was prepared as previously described (9). AKTVIII (Sigma), Imatinib (LC laboratories I-5508) and BGB324/R428, BGB214, were prepared in DMSO. Cell culture, retroviral transductions, siRNA transfection HMEC strains (4th passage) were established and maintained as described (67) in M87A medium with oxytocin and cholera toxin (68). PANC1, MDA-MB-231, 4T1, and MCF10A (American Type Culture Collection, Rockville, MD) cells were cultured as described (11). HMLE and HMLER cells (a gift from Dr. R. Weinberg) were maintained as per Mani et al (69). TBK1 WT and deficient KIC murine cell lines were established and maintained as previously described (14). siRNA transfections were conducted as previously described (70).

### CRISPR Knockout

Oligos of the gRNAs were annealed with T4 polynucleotide kinase (New England Biolabs) by PCR. Annealed oligos were then ligated to PX458 vector with FastDigest BbsI (FD1014, Thermo Fisher Scientific) and T7 DNA Ligase (M0318, New England Biolabs) by PCR. Mixture from the reaction was then transformed into the NEB 5-α Competent *E. coli* (High Efficiency). DNA was extracted from expanded colonies and sent to UTSW sequencing core for sequencing. Plasmids with correct gRNA sequences or empty vector control were transfected into *KPFC* cells with Lipofectamine 2000 (11668027, Thermo Fisher Scientific). Positive cells expressing GFP were sorted as single clones and expanded. Each expanded clone was subjected to validation through PCR and western blot analysis.

Oligos used for the cloning of gRNAs (KO A and KO B) targeting *AKT3* are

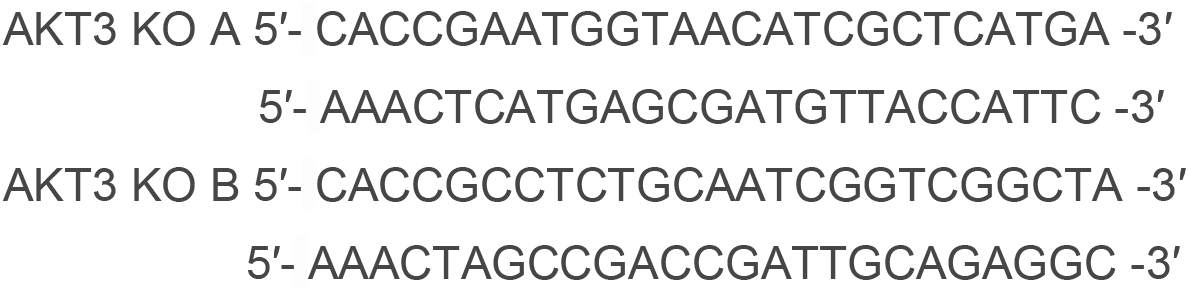

Primers for PCR validation are

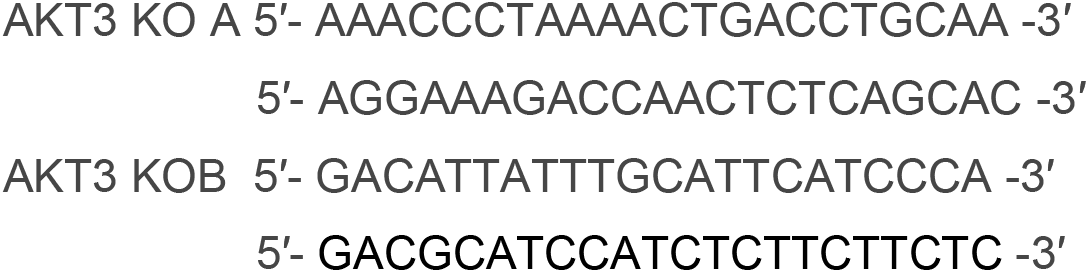

### Immunoblotting and Flow Cytometry

Western blot analysis and flow cytometry analysis of cell lines were conducted as previously described (11). MCF10A cells were treated with TGFβ (10 ng/ml) for 4 days and then lysed using NP40 Cell Lysis Buffer (40 mM HepesNAOH, 75 mM NaCl, 2 mM EDTA, 1% NP40, phosphatase inhibitor cocktail tablet, protease inhibitor cocktail tablet (Roche)). For immunoprecipitation, antibodies against separate AKT isoforms (1, 2 and 3) and control IgG antibodies (1μg/lysate) were added to lysates and incubated overnight at 4°C. Next day the pre-blocked protein-A/G beads (GE Healthcare) in lysis buffer were added and allowed to bind at 4°C for 1 hr. Beads were then washed 3 times (20 mM Tris-HCl (pH 7,5), 150 mM NaCl, 1% NP40) and protein eluted by boiling in SDS-PAGE loading buffer. Running of SDS/PAGE gel and immunoblotting were carried out according to standard procedures. Membranes were probed using anti-pAKT (Ser473) and Pan-AKT antibodies. Nuclear extraction of MDA-MB-231 cells was done according to manufactureŕs instructions (Universal Magnetic Co-IP Kit, Active Motif, 54002). Imaging flow cytometry analysis was conducted on an Amnis Imagestream Mk (>100,000 events) using the Imagestream software (Tree Star, Inc., Ashland, OR, USA) in the Flow Cytometry Core at UT Southwestern. All Western and flow cytometry results shown were performed in at least three independent experiments.

### Biochemical assays for AKT activity

An Akt activation assay was used in which PDK1 was used to phosphorylate inactive Akt enzymes, which then phosphorylated a GSK3α-derived LANCE *Ultra* U*light*-labelled crosstide substrate (Perken Elmer, TRF0106-M). Addition of a Europium-labelled antibody specific to the phosphopeptide (LANCE *Ultra* Europium-anti-phospho-Crosstide (anti-GSK-3α Ser21, Perkin Elmer, TRF0202-M) allows proximity-dependent energy transfer from the Europium donor to the *Ultra* U*light*^TM^ acceptor. Briefly, 5 µL enzyme in 1X AB (50 mM HEPES pH 7.5, 1 mM EGTA, 10 mM MgCl2, 0.01% Tween, 2 mM DTT) was incubated with 2.5 µL test compound. To start the reaction 2.5 µL reaction mix was added which consisted of PDK1, lipid preparations, crosstide and ATP in 1X AB. Final assay concentrations were: 1% DMSO, 5 nM Akt1/5-15 nM Akt2/3-5 nM Akt3 as appropriate, 5 nM PDK1, 5.5 µM DOPS, 5.5 µM DOPC, 0.55 µM PtdIns(3,4,5)P3, 100 µM ATP, 100 nM crosstide. After 30 min, the reaction was stopped using 5 µL 40 µM EDTA in 1X LANCE Detection buffer (Perkin Elmer, CR97-100) for 5 min. For detection, 5 µL 8 nM Europium-anti-phospho-Crosstide antibody in 1X Detection buffer was added to each well and incubated for 1 h. Plates were read with an EnVision® Multilabel Plate Reader, excitation at 320 nm and emission at 665 nm and 615 nm. Results were converted to percent inhibition of phosphorylation by normalizing to positive and negative controls, and compound IC50 was determined using a 3-parameter equation (Prism, GraphPad).

### 3D culture experiments

Growth factor reduced Matrigel (Corning, 10–12 mg/mL stock concentration, catalog no. 354230) and bovine (Corning, catalog no. 354231) or rat tail (Corning, catalog no. 354236) Collagen I were used for organotypic culture experiments. Vertical invasion assays and experiments in three-dimensional (3D) culture were performed and quantified as described previously (71) using a Matrigel/Collagen I matrix (3–5 mg/mL Matrigel and 1.8–2.1 mg/mL Collagen I). A 120-μm span on the *z*-axis is shown for the vertical invasion assays.

### Mammosphere and tumorsphere formation assay

Mammosphere cultures were performed as previously described (72). Single cells were plated in ultra-low attachment plates (Corning, Acton, MA, USA) at a density of 20,000 viable cells/ml. Total mammospheres per well were quantified using ImageJ.

### Gene Expression Analysis and RNA sequencing

The expression analysis of the breast cancer cell lines and human samples (cancer, normal) was performed from published and GEO-submitted Affymetrix data as described (Kilpinen et al., 2008). Global gene expression analysis of HMEC lineage was performed on FACS sorted (FACSVantageSE) pre-stasis HMEC (4th passage) cells. Total RNA from FACS-enriched primary culture cells were isolated with TRIzol (Invitrogen) and RNeasy Mini column (Qiagen) and evaluated using Bioanalyzer (Agilent Technologies). Gene expression levels were measured using the Illumina HumanHT-12 v4 Expression BeadChip whole-genome expression array. The Illumina Bead Array data were quality controlled in Genome Studio and both probe level and gene level data were imported into JExpress Pro (http://jexpress.bioinfo.no) for analysis. After quantile normalization both datasets were log2 transformed. Correspondence Analysis (Fellenberg et al., 2001) was performed on the datasets, together with Hierarchical Clustering of the samples using a Pearson correlation measure on a per gene mean centered version of the data. Differentially expressed genes between AXL^+^ and AXL^-^ groups were identified using the Rank Product method on both datasets (Breitling et al., 2004). The resulting lists of differentially expressed genes with a false discovery rate value q=10% from these two analysis was considered differentially expressed between the two groups. Cells were plated on 10 cm dishes until cell densities of 70% were achieved. Total RNA was extracted from cells using QIAGEN RNeasy Mini kit and stored at -80°C. 1 μg total RNA per sample were subjected to library generation using the TruSeq stranded total RNA sample preparation kit, according to the manufacturer’s protocol (Illumina). The libraries were pooled and sequenced on a NextSeq 500 instrument (high output flowcell) at 1x75 bp single end reads (Illumina). Raw RNAseq reads were aligned against to the human genome release GRCh38/hg38 using HISAT2 (73) and exons were counted using RSubread.featureCounts (74). Libraries were filtered to remove gene counts of less than 1 CPM across all libraries and normalized. Differentially expressed genes between GFP control group and AKT3 overexpressing MCF10A cells were calculated using edgeR (75, 76). Genes were considered differentially expressed with a fold change >2 and p<0.05.

### AKT3 score and Metabric dataset

To assess the influence of AKT3 signaling and its downstream targets on survival of breast cancer patients, genes that were found to be differentially expressed after AKT3 overexpression in MCF10A cells were used to generate an AKT3 score. The score essentially represented the sum of expression of 42 differentially expressed genes, adjusted for expected directionality. Initially, we examined 46 different genes, but only 42 of them were represented with probes on the expression array. For genes represented by multiple probes (the 42 genes mapped to 71 different probes), mean signal intensity was used. The influence on breast cancer specific survival and the putative difference between molecular subtypes was investigated in the Metabric cohort, composed of 1980 breast cancer patients enrolled at five different hospitals in the UK and Canada (77). Gene expression was assessed using the Illumina HT-12 v3 microarray and normalized data was downloaded from the European Genome-phenome Archive (EGA) data portal. Missing values were imputed using the impute.knn function as implemented in the R library ‘impute’ with default settings (Hastie T, c R, Narasimhan B and Chu G (2016). Impute: Imputation for microarray data. R package). The data was batch adjusted for hospital effect using the pamr.batchadjust function in the ‘pamr’ library with default settings (T. Hastie, R. Tibshirani, Balasubramanian Narasimhan and Gil Chu (2014). Pam: prediction analysis for microarrays). Association between the score and molecular subtypes (77, 78) was tested using Kruskal-Wallis rank test, and correlations were estimated with Spearman’s rank correlation. Survival analyses were performed using Cox proportional hazards regression model as implemented in the R library ‘rms’ (Frank E Harrell Jr (2016). rms: Regression Modeling Strategies). Survival plots were generated using the survplot function, as implemented in the rms library. All analyses were performed using R version 3.3.1.

### Confocal Microscopy

Cells were plated on coverslips (79.5, Marienfeld-Superior) overnight under low serum (1%) conditions. Cells were fixed with 4% formaldehyde diluted in warm PBS for 15 min, washed 3 times, blocked, and permeabilized with 5% goat serum, 0.3% Triton X100 in PBS for 1h. Cells were incubated with the appropriate primary antibody overnight followed by 3 wash steps with PBS and secondary antibody incubation for 2h in 5% BSA in PBS. After 3 wash steps with PBS, coverslips were mounted on slides with Prolong Diamond Antifade Reagent (Thermo Fisher). The images were acquired using Leica SP5, Leica SP8, Zeiss LSM780, or Zeiss LSM880 inverted microscopes.

### Immunohistochemistry

Paraffin-embedded Human PDAC samples were provided by the Tissue Management Shared Resource within the Simmons Comprehensive Cancer Center at UT Southwestern. Both AXL and AKT3 antibodies were optimized and stained using a Leica Autostainer. Paraffin-embedded normal human breast tissue sections (n=20; generously provided by Dr. A.Borowsky) were prepared for immunofluorescence and stained with as previously described (Garbe et al., 2012). For N-cadherin analysis, antigen retrieval was performed by boiling for 20 min at in Tris EDTA buffer, ph 9 in a microwave oven. A Dako Autostainer was used for staining. The slides were incubated 60 minutes at room temperature with a monoclonal antibody against N-cadherin (M3613), dilution 1:25 (Dako). Immunoperoxidase staining was carried out using the Dako Envision Kit with diaminobenzidin tetrachloride peroxidase. For analysis of Twist-2, antigen retrieval was performed by boiling in TRS buffer (pH 6.0) (Dako) for 25 minutes, and incubated for 1 hr in room temperature with the rabbit polyclonal antibody Twist-2 diluted 1:500, and stained with HRP EnVision rabbit (Dako) for 30 minutes in RT. The peroxidase was localized by the diaminobenzidine tetrachloride peroxidase reaction and counterstained with Mayer’s hematoxylin. For Axl analysis, the sections were boiled in TRS buffer (pH 6.0) (Dako) in 20 minutes, followed by incubation overnight at room temperature with goat IgG antibody Axl, dilution 1:50 (R&D AF854) and stained with EV rabbit for 30 minutes. The peroxidase was localized by the diaminobenzidine tetrachloride peroxidase reaction and counterstained with Mayer’s hematoxylin. The human breast cancer tumor sections were obtained from the IRO database and assayed for quality control by a pathologist. IHC staining was carried out using DAKO, EnVision™ FLEX kit with DAB before counterstaining with hematoxylin (DAKO, EnVision™ FLEX Hematoxylin K8008). Stained samples were acquired using with Zeiss Axio Observer Z1 microscope and analyzed with TissueGnostics software for acquisition and analysis. Representative regions were analyzed from each sample slide and mean intensity of DAB-AKT3 staining from nuclei and cytoplasm was used to separate nuclear AKT3 cases from cytoplasmic AKT3 cases.

### Animal studies

#### Sygeneic pancreatic cancer model

*KPFC* (CAS9-EV, AKT3 KO, Rescue) cells were injected orthotopically (2.5 × 10^5^ cells) in 6-to 8-week-old C57BL/6 mice. 19 days after tumor cell injection mice were sacrificed and organs were harvested for analysis. All animals were housed in a pathogen-free facility with 24-h access to food and water. Animal experiments in this study were approved by and performed in accordance with the institutional animal care and use committee at the UTSW Medical Center at Dallas. Before implantation, cells were confirmed to be pathogen free.

#### Tumor cell titration studies

Xenograft tumor-initiation studies were conducted as described by (80). HMLER cells (GFP, myrAKT1, myrAKT3 or AKT3) were suspended in DMEM/Matrigel (1:1) in 50 μL) and injected subcutaneously into 3-6 weeks old NOD-SCID mice. Tumor incidence was monitored with hand held caliper for up to 60 days after injection; tumor threshold was set at 20 mm3 (AKT3) or 25 mm3 (myrAKT3). Animals were treated with BGB214 dissolved in 0.5% HPMC/0.1% Tween 80 (Vehicle) as indicated in figure legends starting at the day of cell injection. For some studies, cells were treated for 24 hrs with 0.54 μM BGB214 prior to implantation. Tumor cell titration animal experiments were approved by The Norwegian Animal Research Authority andperformed in accordance with The European Convention for the Protection of Vertebrates Used for Scientific Purposes.

### In vitro kinase assay

1.5 μg of recombinant GST-AKT3 (BML-SE369-0005, Enzo Life Sciences), 0.1 μg of Recombinant Active TBK1 (T02-10G-05, SignalChem), and 1 μL 10 mM ATP were combined in kinase reaction buffer (20 mmol/L Tris-HCl pH 7.4, 500 mmol/L β-glycerol phosphate, 12 mmol/L magnesium acetate) up to a total of 30 μL. Kinase reaction was carried out at 30°C, 500 rpm for 1hr. After reaction, AKT3 protein was resolved by SDS-PAGE and stained by Coomassie Brilliant Blue. Bands were cut out for MS analysis to identify phosphorylation site by the UT Southwestern Proteomics core.

### Statistical analysis

GraphPad Prism 5.0 for PC, and MatLAB were used for statistical analysis using tests stated in the Figure Legends. Comparisons of histological SI groups were performed by Pearson X^2^ test using cut-off values for staining index (SI) categories based on median values. Grouped analyses were performed with Bonferroni’s test for multiple comparisons. Significance was established when p<0.05.

## Author contributions

ENA, JBL, and RAB conceived and designed the study. ENA, JMW, SH, CET, MB, AM, AV, NP, SN, AR, JET, TR, VF, DM, and KYA acquired data and performed analysis and interpretation of data. ENA wrote the manuscript. RAB and JBL reviewed and revised the manuscript. GG, JI, JBL, and RAB supervised the study.

## Acknowledgements

We thank Sissel Vik Berge, Karla Sputova, Edward Verwayen, Hallvard Haugen, Marianne Enger, Michelle Scottn, Anna Boniecka, and Eline Milde for excellent technical support, and to Dr. Kjell Petersen, the Computational Biology Unit, the University of Bergen, and the Norwegian Bioinformatics platform for microarray analysis support. Additionally, we thank our UT Southwestern colleagues Lianxin Hu and Qing Zhang for their input and technical advice. We would also like to thank the UT Southwestern Proteomics core, Cheryl Lewis, director of the Shared Tissue Management Resource Core (supported by P30 CA142543), and Angie Mobley, manager of the Flow Cytometry core. The work was supported by NIH grants R01 CA192381, R01 CA243577 and U54 CA210181 Project 2 to RAB, the Effie Marie Cain Scholarship in Angiogenesis Research and the Gillson Longenbaugh Foundation to RAB, grants to JBL from the Norwegian Cancer Society, Norwegian Research Council, Bergen Health Authority and BerGenBio ASA, as well as CPRIT RP160157 (principal investigator: M. Cobb, UT Southwestern, Dallas, Texas) and NCI F99 CA253718 to ENA. The funders had no role in study design, data collection and analysis, decision to publish, or preparation of the manuscript.

