## Supplemental Figures and Tables for "AXL-TBK1 driven nuclear AKT3 promotes metastasis"

**Number of Supplementary figures, tables:** 6 supplemental, 1 supplemental table

**Supplemental Figure 1.** AXL signaling is required for EMT in breast cancer

**Supplemental Figure 2.** AKT3 is associated with EMT

**Supplemental Figure 3.** EMT requires AXL dependent activation of nuclear localized EMT

**Supplemental Figure 4.** Nuclear Localization of AKT3 is dependent on NLS sequence

**Supplemental Figure 5.** AKT3 correlates with slug in invasive breast carcinoma

**Supplemental Figure 6.** Evaluation of BGB214 in vitro and in vivo

**Supplemental Table 1.** Description of cell lines used in study

**Supplemental Table 2.** Differentially expressed (DE) genes with AKT3 overexpression

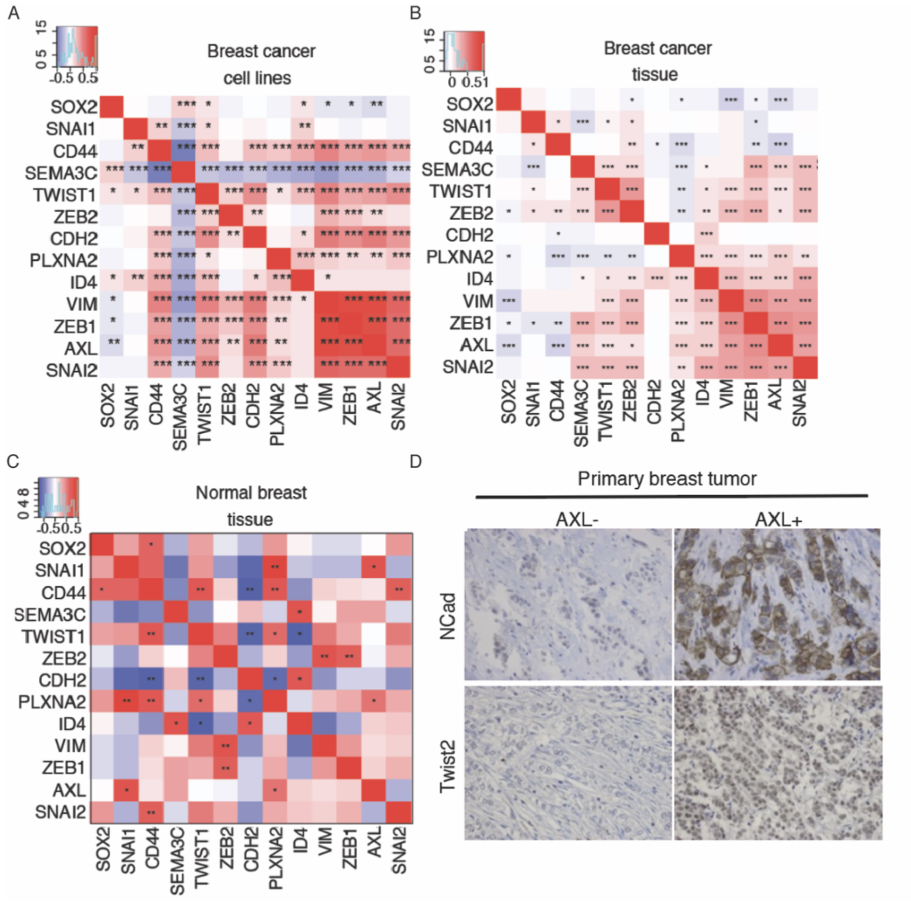


**Supplemental Figure 1. AXL signaling is required for EMT in breast cancer.**

**A-C**) *AXL* mRNA correlates with EMT and stem cell-related gene expression in breast cancer cell lines (**A**), breast carcinoma biopsies (**B**) and normal breast tissue (**C**). Positive correlation values are demarcated as red and negative correlation values are shown as blue (* p<0.045, ** p<0.009, *** p <2x10-5; Spearman's correlation test). **D**) IHC of patient primary breast tumor biopsies with strong AXL expression correlate with EMT marker N-cadherin and twist2 expression.

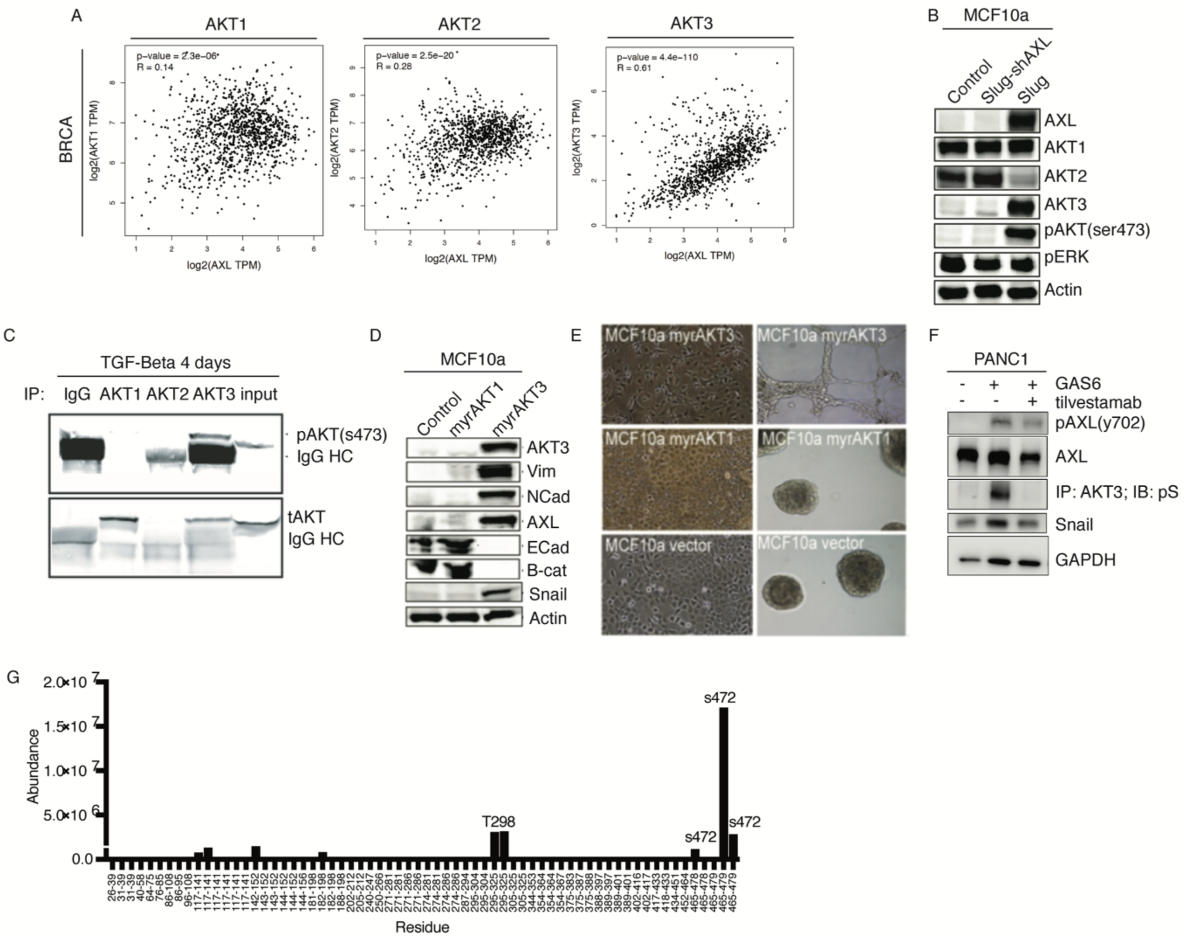


**Supplemental Figure 2. AKT3 is associated with EMT and breast tumor initiation**

**A**) Gene expression correlation of AKT isoforms and AXL using GEPIA (Gene expression profiling interactive analysis) in invasive breast carcinoma (BRCA) and pancreatic adenocarcinoma (PAAD) tumors from the TCGA and GTEx databases. **B**) Protein level of AKT isoforms (AKT-1,-2,-3), Axl, total phosphorylated AKT S473 (pAKT) and ERK1/2 (pERK) in MCF10a -expressing slug, or control vector, and MCF10a/slug transduced with shAXL or shLuc retroviral vector. **C**) Immunoprecipitation of AKT isoforms (AKT-1,-2,-3) from MCF10a cell extracts after TGFβ-treatment (10 ng/ml, 4 days) followed by Western blot for phospho-AKT S473 show that AKT3 is the main phosphorylated AKT isoform. **D**) MCF10a cells were transduced with retroviral vectors that express myrAKT1 or myrAKT3 and analyzed for AXL, epithelial (E-cadherin) and mesenchymal (vimentin, N-cadherin) marker expression, AKT1/3 and pAKT levels. **E**) Phase contrast images of MCF10a cells in 2D tissue culture plates (left) and matrigel (right). **F**) PANC1 cells were stimulated with DMSO, GAS6 (200 ng/ml) +/- anti-AXL tilvestamab. Immunoprecipitation of AKT3 was probed for total phosphorylated serine. Total lysates were probed for pAXL y702, total AXL, snail, and GAPDH (loading control). **G**) In-vitro kinase activity assay of human recombinant TBK1 and AKT3 using cold ATP. PTM mass-spec analysis revealed that TBK1 phosphorylates AKT3 at serine 472. Residue abundance of recombinant AKT3 alone was subtracted from residue abundance of activated AKT3. All representative results shown (B-F) were reproduced in at least three independent experiments.

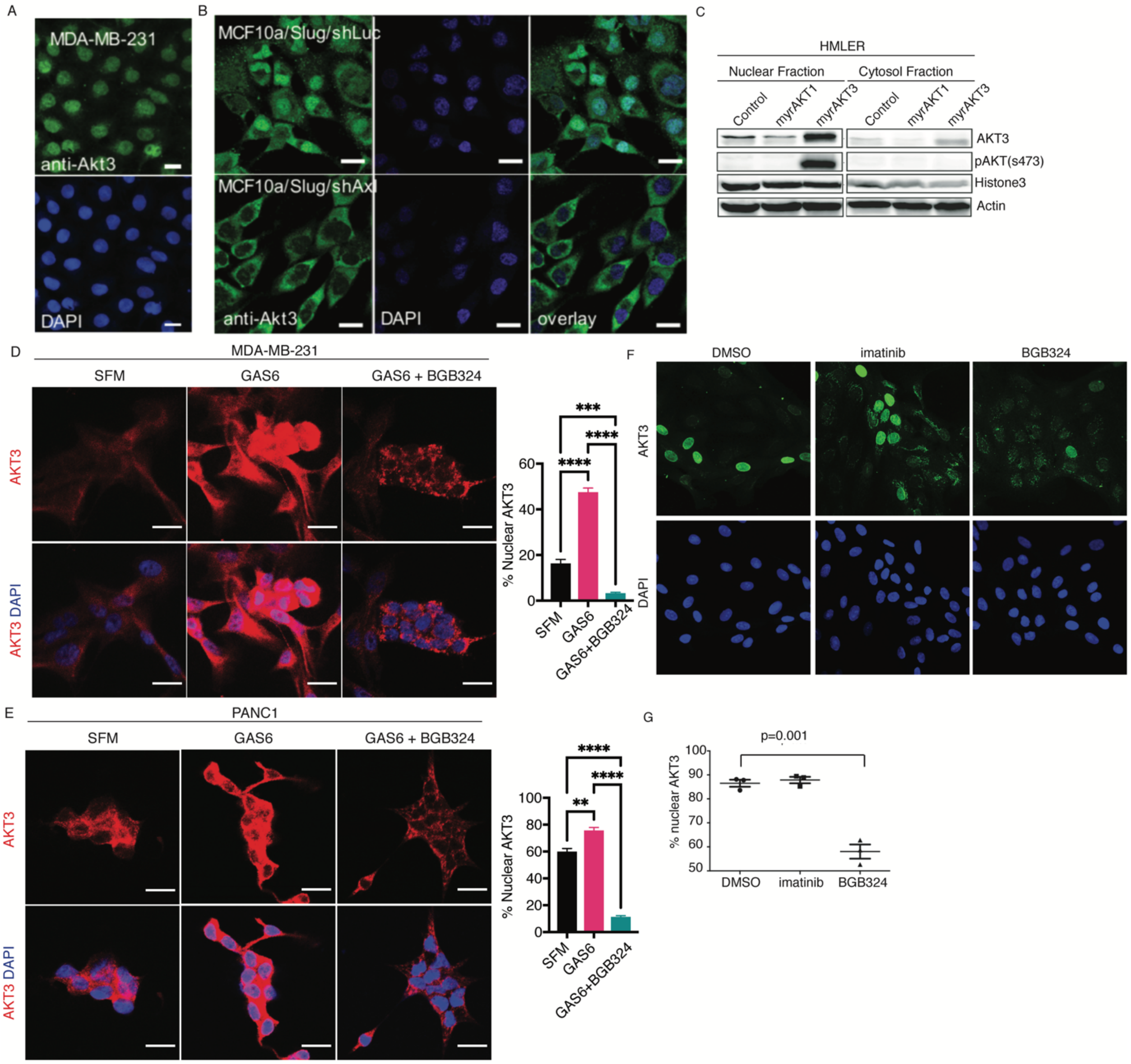


**Supplemental Figure 3. EMT requires AXL dependent activation of nuclear localized AKT3. A**) Subcellular localization of endogenous AKT3 (anti-AKT3-FITC, green) in MDA-MB231 by immunofluorescence. Nucleus: DAPI (blue). Scale bar, 50 μm. **B**) Immunofluorescence analysis of AKT3 (anti-AKT3-FITC, green) MCF10a/slug cells after transfection with shAXL or shLuc retroviral vectors. Scale bar, 20 μm. **C**) Western blot analysis of AKT1 and AKT3, pAKT levels and nuclear marker protein histone 3 in nuclear and cytoplasmic cell fractions from HMLER cells transduced with retroviral vectors encoding myrAKT1, myrAKT3 or control vector. **D-E)** Immunofluorescence of TBK1 (green), AKT3 (Red), and DAPI (blue) in MDA-MB-231 (**D**) and PANC1 (**E**) cells treated with SFM, 200 ng/mL GAS6 +/- 2 μM BGB324 for 12 hrs. Cells were imaged at 20X using confocal microscopy. Scale bar, 20 μm. Percent of cells with nuclear

AKT3 are graphed to the right of each cell type, n>200 cells. Statistics were done use one-way ANOVA; ** p <0.01, *** p <0.001, **** p <0.0001 **F, G**) Nuclear localization (**F**) and quantification (**G**) of AKT3 in AXL-expressing mammary epithelial progenitor cells requires AXL but not ABL/CKIT/PDGFR activity. cKit+ AXL+ HMEC cells treated with vehicle (DMSO), 1 μM imatinib, or 600 nM BGB324 for 24 hrs and analyzed by immunofluorescence (anti-AKT3-FITC, green). Nucleus: DAPI (blue). Scale bar, 50 μm. %nuclear AKT3 shown as mean ± S.E.M. from 3 x 25,000 cells/coverslip, in triplicate; p=0.001, t-test. All representative results shown were reproduced in at least three independent experiments.

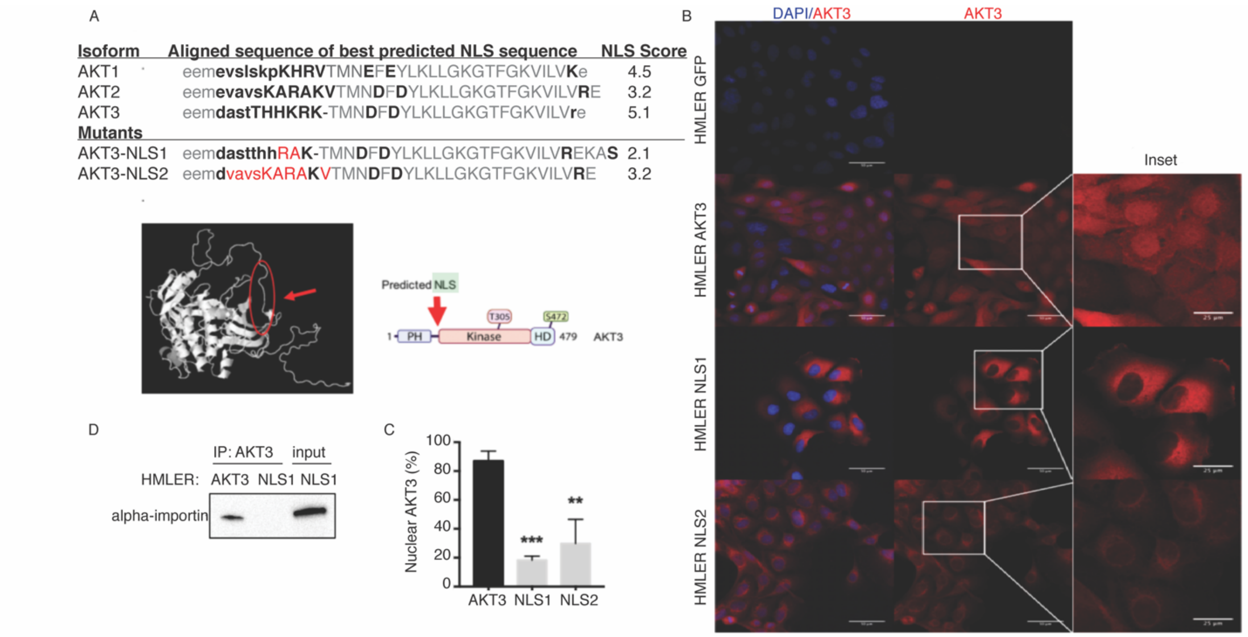


**Supplemental Figure 4. Nuclear Localization of AKT3 is dependent on NLS sequence**

**A**) Display of predicted nuclear localization sequence (NLS) in AKT3 with calculated in silico NLS score as defined in (31). The *in silico* predicted bipartite NLS is depicted in upper case, with flanking residues in lower case. Exact matches between all three isoforms are highlighted in grey, with mutated amino acids in red. The score is based on the predominate staining pattern from 1 (cytoplasmic only), 4 (mixed cytoplasmic and nuclear, mostly cytoplasmic), 7 (mixed cytoplasmic and nuclear, mostly nuclear) to 10 (nuclear only). A score of 5 is the lowest score where the protein is exclusively or primarily in the nucleus in a significant proportion of the cells, although the majority of the cells show mixed nuclear and cytoplasmic staining. **B**) Confocal images of HMLER cells that overexpress AKT3 and AKT3-NLS mutants (red) with quantitation of subcellular location shown in (**C**). At least 60 cells per condition were analyzed. Data shown as percent of cells with predominant nuclear AKT3. **D**) Immunoprecipitation using anti-AKT3 of HMLER-AKT3 and AKT3-NLS1 lysate and blotting for α-importin (lane 1-2). Whole lysate of HMLER AKT3-NLS1 as expression control (lane 3). All representative results shown were reproduced in at least three independent experiments.

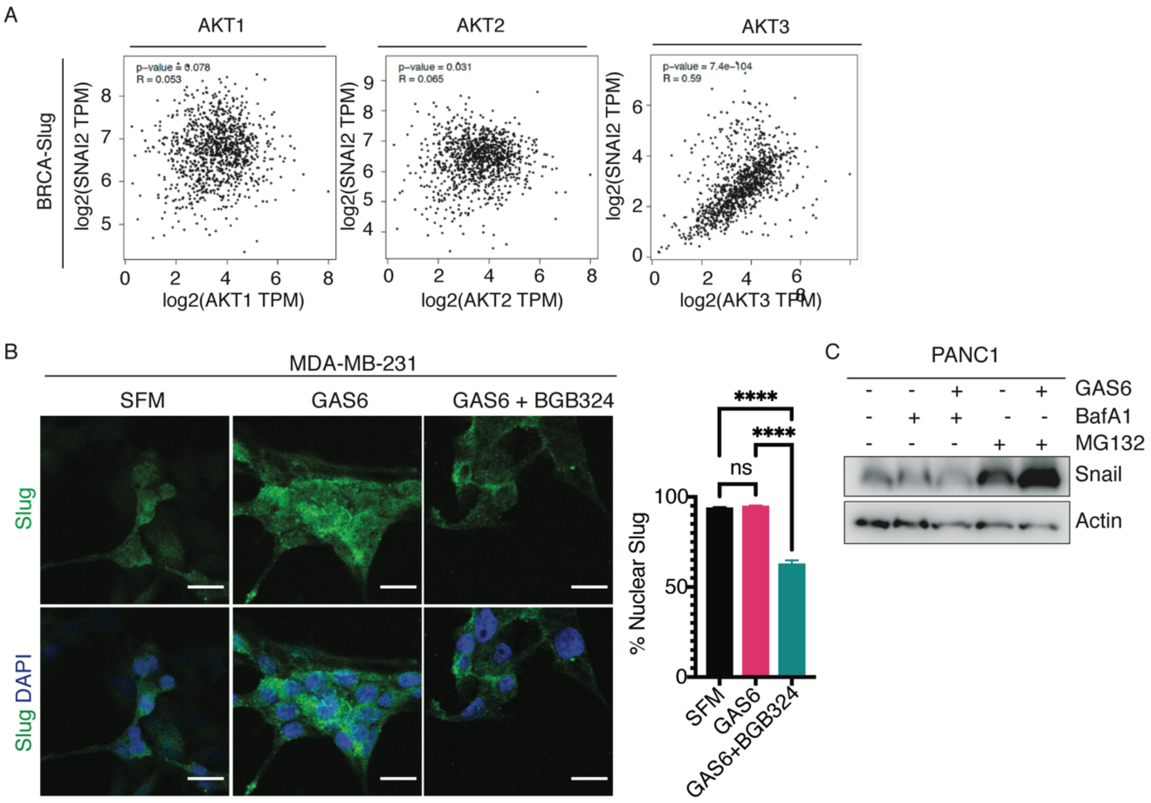


**Supplemental Figure 5. AKT3 correlates with slug in invasive breast carcinoma**

**A)** Gene expression correlation of *AKT* isoforms and *SNAI2* using GEPIA (Gene expression profiling interactive analysis) in invasive breast carcinoma tumors (BRCA) from the TCGA and GTEx databases. **B)** Immunofluorescence of slug (green) and DAPI (blue) in MDA-MB-231 cells treated with SFM, 200 ng/mL GAS6 +/- 2 μM BGB324 for 12 hrs. Cells were imaged at 20X using confocal microscopy (scale bar, 20 μm) and nuclear slug was quantified, n>200 cells. **C)** PANC1 cells were treated for 8 hrs with DMSO, BafA1 (0.5 μM) +/- GAS6 or MG-132 (10 μM) +/- GAS6. Cells were lysed and probed for snail and actin (loading control). Nuclear slug statistics were done use one-way ANOVA; **** $p < 0.0001$.

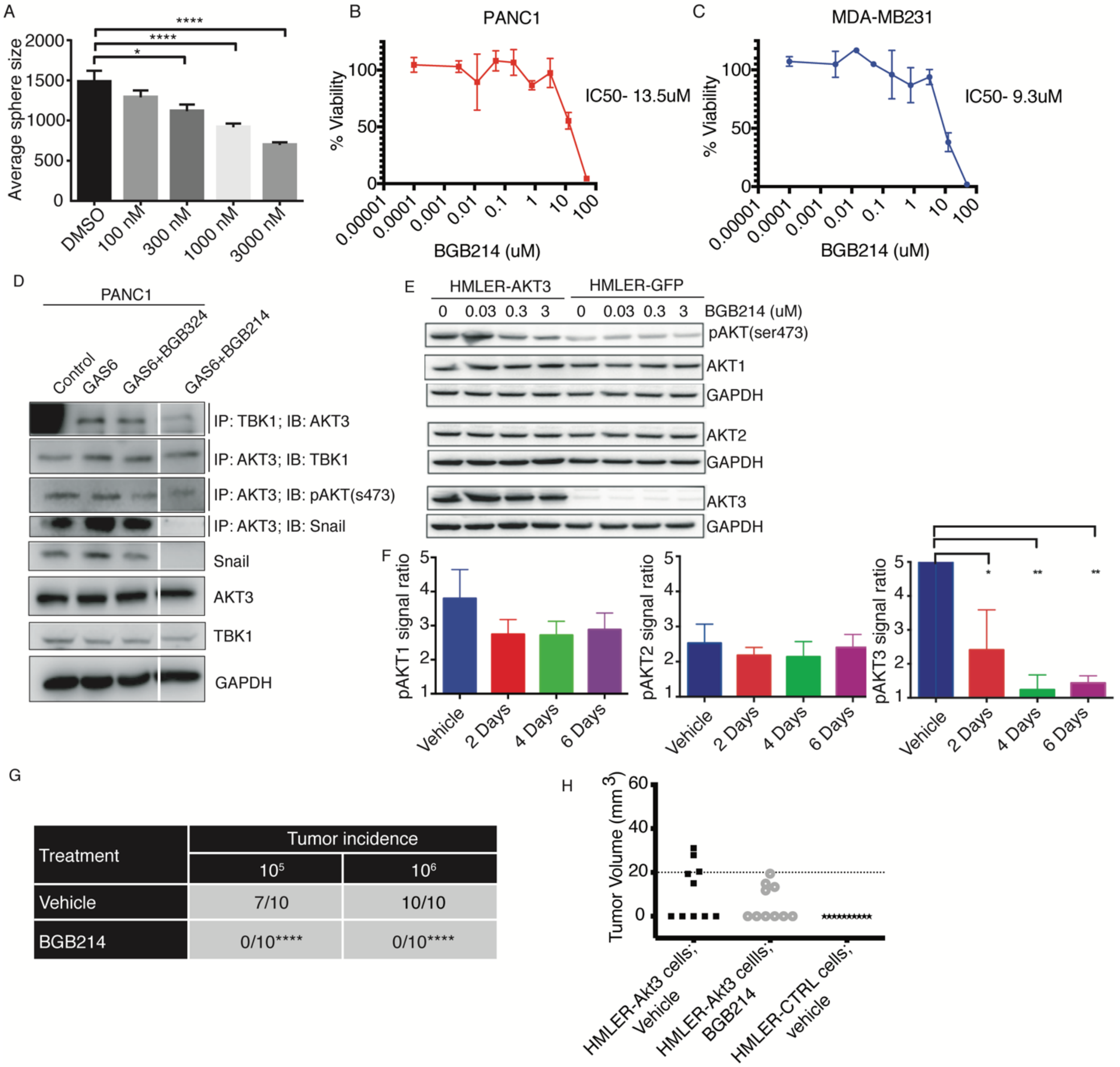


**Supplemental Figure 6. Evaluation of BGB214 in vitro and in vivo**

**A**) MDA-MB-231 cells were seeded in 3D Matrigel cultures and treated for 9 days with the indicated dose of BGB214. Spheres were quantified. Histograms show average sphere size for each treatment. Mean + SE is shown, n=3. Statistical analysis by one-way ANOVA, Tukey's multiple comparisons test. *p < 0.05, ****p<0.0001. **B**) PANC1 (**C**) or MDA-MB-231 cells were plated on day 0 and BGB214 was added on day 1 in 4-fold dilutions. For each assay, 8 different drug concentrations were tested with 8 replicates per concentration. Relative cell number was determined by adding MTS incubating for 1 to 3 hrs at 37°C. Drug sensitivity curves and IC50s were calculated using in-house software. Response was validated in replicate plates (n ≥ 4). Assays have been repeated in 4 biological replicates, with 16 technical replicates each in total.

**D**) PANC1 cells were stimulated with DMSO, GAS6 (200 ng/mL) +/- 2 μM BGB324 or 3 μM BGB214. Immunoprecipitation of AKT3 was probed for pAKT(s473), TBK1, and snail. Immunoprecipitation of TBK1 was probed for AKT3. Lysates were probed for AKT3, snail, TBK1 and GAPDH (loading control). Lanes were run on same blot but space indicates they are not continuous. **E**) HMLER-AKT3 or HMLER-GFP cells were treated with increasing concentration of BGB214 (AKT3i). Lysates were harvested and probed for pAKT(s473), AKT1, -2, or -3. GAPDH was used as a loading control. **F**) MCF10A-DCIS cells were transplanted subcutaneously into mice, treated with 25 mg/kg BGB214 for 2-6 days. pAKT1, 2, and 3 levels were subsequently evaluated. **G**) Tumor incidence of HMLER/IRES-GFP and HMLER/Akt3 cells injected subcutaneously into host NSG mice at limiting dilutions (between $10^5$-$10^6$ cells) followed by treatment with BGB214 (150 mg/kg, orally once daily in a 5 days on 2 days off regimen). Cells were treated in vitro for 24 hrs with BGB214 at 0.5 μM. Tumor incidence was evaluated 14 days post implantation and tumor size cut off value was set at 20 $mm^3$. **H**) Tumor incidence of HMLER/IRES-GFP and HMLER/Akt3 cells injected subcutaneously into host NSG mice at limiting dilution ($10^6$ cells) and treated with BGB214 (50 mg/kg, orally twice daily). Tumor incidence was evaluated 14 post implantation and tumor size cut off value of 20 $mm^3$ is indicated. **** p 0.0001 by unpaired Student two tailed test

| Cell Name | Cell Type | Species | AXL | AKT3 | snail | slug | Source |
|---|---|---|---|---|---|---|---|
| **KPFC** | Cancer; Pancreatic ductal adenocarcinoma; primary cell line; KRAS G12D; P53 deletion | mouse | + | + | - | + | (80) |
| **KIC** | Cancer; Pancreatic ductal adenocarcinoma, primary cell line; KRAS G12D; INK4A deletion | mouse | + | + | + | + | (14) |
| **PANC1** | Cancer; Pancreatic ductal adenocarcinoma | human | + | + | + | - | (81) |
| **MDA-MB-231** | Cancer; Triple negative breast adenocarcinoma | human | + | + | - | + | (82-84) |
| **MCF10A** | Epithelial mammary gland; Fibrocystic disease | human | - | - | - | - | (85, 86) |
| **MCF10A/slug** | Epithelial normal breast cells transduced with slug | human | - | - | - | + | (87) |
| **HMEC** | Primary mammary epithelial cells | human | - | - | - | + | (88, 89) |
| **HMLER** | HMEC cells transformed by transduction with hTERT, SV40LT, and RAS oncogenes | human | - | - | - | - | (90) |

**Supplemental Table 1**. Description of cell lines used in study. Protein expression of AXL, AKT3, snail, and slug is listed for each cell line used in study.

| Symbol | logFC | Pvalue | FDR |
|---|---|---|---|
| **AKT3** | 4.96 | 1.40E-27 | 2.12E-23 |
| **TGFBI** | 1.78 | 4.53E-15 | 2.30E-11 |
| **CD24** | 1.89 | 2.91E-14 | 1.10E-10 |
| **SERPINE1** | 1.6 | 5.67E-12 | 1.72E-08 |
| **A2ML1** | 1.49 | 1.18E-11 | 3.00E-08 |
| **SPRR1B** | 1.73 | 2.77E-11 | 6.01E-08 |
| **KLHL4** | 4.26 | 8.05E-11 | 1.53E-07 |
| **IGFBP3** | 1.84 | 1.05E-10 | 1.78E-07 |
| **ZBED2** | 1.91 | 1.50E-10 | 2.28E-07 |
| **OVOL1** | 2.51 | 7.09E-10 | 9.80E-07 |
| **KRT6A** | 1.31 | 9.94E-10 | 1.26E-06 |
| **F3** | 1.66 | 1.51E-09 | 1.77E-06 |
| **IL1A** | 1.84 | 2.34E-09 | 2.54E-06 |
| **IVL** | 2.3 | 1.07E-08 | 1.09E-05 |
| **IL7R** | 1.95 | 2.24E-08 | 2.13E-05 |
| **SDK2** | -5.3 | 4.58E-08 | 4.09E-05 |
| **S100P** | 1.41 | 6.26E-08 | 5.28E-05 |
| **PI3** | 1.44 | 7.71E-08 | 6.16E-05 |
| **LAMB3** | 1.22 | 1.24E-07 | 9.13E-05 |
| **UCA1** | 2.3 | 3.47E-07 | 2.39E-04 |
| **KRT6C** | 1.8 | 4.26E-07 | 2.70E-04 |
| **LAMC2** | 1.24 | 4.19E-07 | 2.70E-04 |
| **CPA4** | 1.4 | 8.50E-07 | 5.16E-04 |
| **WNT7B** | 1.55 | 1.33E-06 | 7.77E-04 |
| **SPINK6** | 2 | 1.60E-06 | 9.00E-04 |
| **CEACAM5** | 3.29 | 1.67E-06 | 9.06E-04 |
| **SERPINB2** | 2.59 | 2.19E-06 | 1.11E-03 |
| **LTBP1** | 1.45 | 5.43E-06 | 2.66E-03 |
| **ANGPTL4** | 1.21 | 6.17E-06 | 2.93E-03 |
| **IL1B** | 1.75 | 7.20E-06 | 3.31E-03 |
| **ADGRF4** | 1.58 | 7.64E-06 | 3.41E-03 |
| **NA** | 1.27 | 7.97E-06 | 3.46E-03 |
| **RYR2** | 1.45 | 1.35E-05 | 5.56E-03 |
| **INHBA** | 1.32 | 1.72E-05 | 6.90E-03 |
| **SPRR2D** | 1.52 | 2.04E-05 | 7.93E-03 |
| **KCNJ15** | 3.29 | 2.38E-05 | 9.04E-03 |
| **BMP2** | 1.69 | 2.69E-05 | 9.95E-03 |
| **AC007879.7** | 1.39 | 3.19E-05 | 1.15E-02 |
| **SERPINB10** | 3.01 | 4.16E-05 | 1.47E-02 |
| **SERPINB13** | 1.41 | 4.36E-05 | 1.51E-02 |
| **ZNF114** | 2.33 | 4.59E-05 | 1.55E-02 |
| **PTHLH** | 1.38 | 5.36E-05 | 1.77E-02 |
| **MYLK** | 1.4 | 6.61E-05 | 2.09E-02 |
| **PCDH7** | 1.59 | 6.95E-05 | 2.15E-02 |
| **PSCA** | 1.25 | 1.11E-04 | 3.38E-02 |

**Supplemental Table 2.** Differentially expressed (DE) genes determined by RNA sequencing in MCF10A cells after AKT3 overexpression compared to GFP control vector (DE genes with fold change ≥2, and FDR<0.05).
